# Dynamic regulation of single-cell contractile heterogeneity in the human trabecular meshwork

**DOI:** 10.64898/2026.11.06.764908

**Authors:** Luis Sanchez, Andrew C. Jubintoro, Ivan Pushkarsky, Robert Damoiseaux, Dino Di Carlo, Jie J. Zheng

**Affiliations:** Jules Stein Eye Institute, Department of Ophthalmology, David Geffen School of Medicine, University of California, Los Angeles, CA, USA; Molecular Biology Institute, University of California, Los Angeles, CA, USA; Department of Bioengineering, University of California, Los Angeles, Los Angeles, CA, USA; Department of Molecular and Medicinal Pharmacology, University of California, Los Angeles, Los Angeles, CA, USA; California NanoSystems Institute, University of California, Los Angeles, Los Angeles, CA

## Abstract

Altered trabecular meshwork (TM) mechanics contribute to increased aqueous humor outflow resistance and elevated intraocular pressure in glaucoma, yet how contractility is regulated across individual TM cells remains poorly understood. Using a high-throughput single-cell contractility assay, we quantified time-resolved contractile behavior in primary human TM cells under defined extracellular matrix (ECM) compositions, substrate stiffnesses, and pharmacologic perturbations, and related functional mechanical behavior to intracellular protein abundance. Single-cell analyses revealed that TM contractility is not a uniform population property but a context-dependent contractile state shaped by both extracellular and intracellular regulation. ECM composition established distinct contractile programs, with collagen IV promoting rapid convergence toward a highly contractile state, collagen I supporting a slower but similar response, and fibronectin maintaining a persistently heterogeneous low-contractility state. In contrast, the intracellular perturbation dexamethasone (DEX) did not uniformly increase contractility but selectively prolonged contractile activity within the most mechanically active cells, thereby sustaining highly contractile states. This response was progressively constrained as ECM stiffness increased and was absent on pathologically stiff substrates, demonstrating that extracellular mechanical context conditions glucocorticoid responsiveness. Conversely, ROCK inhibition overrode both basal and DEX-induced contractile programs, collapsing the heterogeneous population into a uniformly low-contractility state. Finally, intracellular protein abundance poorly predicted functional mechanical behavior, with weak pooled associations reflecting divergent donor-specific relationships rather than consistent molecular-mechanical coupling. Together, these findings show that TM contractility emerges from integrating intrinsic cellular state and the extracellular mechanical environment rather than from either alone. More broadly, they identify single-cell contractile state as a functional dimension of TM cell identity that cannot be inferred from static molecular marker abundance or population-averaged mechanical measurements.

## Introduction

Glaucoma is the leading cause of global irreversible blindness^1^. Although age, ethnicity, gender, environmental, and genetic factors all contribute to glaucoma susceptibility^2,3^, elevated intraocular pressure (IOP) remains the primary modifiable risk factor targeted to slow disease progression^4^. IOP homeostasis depends on the trabecular meshwork (TM), which regulates aqueous humor outflow resistance through coordinated interactions between resident cells and their extracellular matrix (ECM). The mechanical behavior of TM cells, including their ability to generate and respond to contractile forces, is therefore fundamental to maintaining normal outflow facility and IOP^5,6^.

TM contractility has been investigated using collagen gel contraction assays, hydrogel-based culture systems, and force-based measurements of cells and tissues. These approaches have established that cytoskeletal tension, cell–matrix interactions, and pharmacologic modulation influence TM mechanics and outflow function^7^. However, nearly all existing methods quantify mechanical behavior as a population average, obscuring the organization of contractile function across individual cells. As a result, it remains unknown whether TM contractility represents a uniform property shared by all cells or emerges from heterogeneous and dynamic behaviors within the population. This limitation also restricts understanding of how contractile responses evolve over time and how distinct cellular subpopulations contribute to tissue-level mechanics.

The TM extracellular matrix (ECM) is compositionally heterogeneous, comprising fibrillar and basement membrane components that provide distinct biochemical and mechanical cues^5,8^. ECM identity and stiffness regulate cellular force generation and mechanotransduction, thereby shaping TM cell contractility^9^, Conversely, contractile forces generated by TM cells actively remodel the surrounding ECM and contribute to its mechanical properties, establishing a dynamic reciprocal relationship between cells and their microenvironment. This bidirectional mechanical feedback is thought to play a central role in maintaining aqueous humor outflow homeostasis and is disrupted in glaucoma. Pharmacologic modulators of TM function, including glucocorticoids and Rho-associated kinase (ROCK) inhibitors, further alter cytoskeletal organization and contractility, yet it remains unclear whether these perturbations act uniformly across the TM cell population or selectively influence distinct mechanical subpopulations. Addressing these questions requires approaches capable of resolving dynamic contractile behavior at single-cell resolution.

Here, we quantified time-resolved contractile behavior in individual primary human trabecular meshwork (HTM) cells using a high-throughput single-cell contractility assay based on ECM micropattern deformation (Fig. 1)^10,11^. This platform enabled direct measurement of contractile dynamics across defined ECM compositions, substrate stiffnesses, and pharmacologic perturbations, including dexamethasone (DEX) and the ROCK inhibitor Y-27632. By integrating single-cell measurements of contractile displacement with intracellular protein expression, we further asked whether molecular identity predicts mechanical function. We hypothesized that TM contractility is an emergent, context-dependent phenotype governed by ECM composition and mechanical environment, and that pharmacologic perturbations reshape the distribution of contractile states rather than uniformly altering the behavior of all cells.

**Figure 1.**
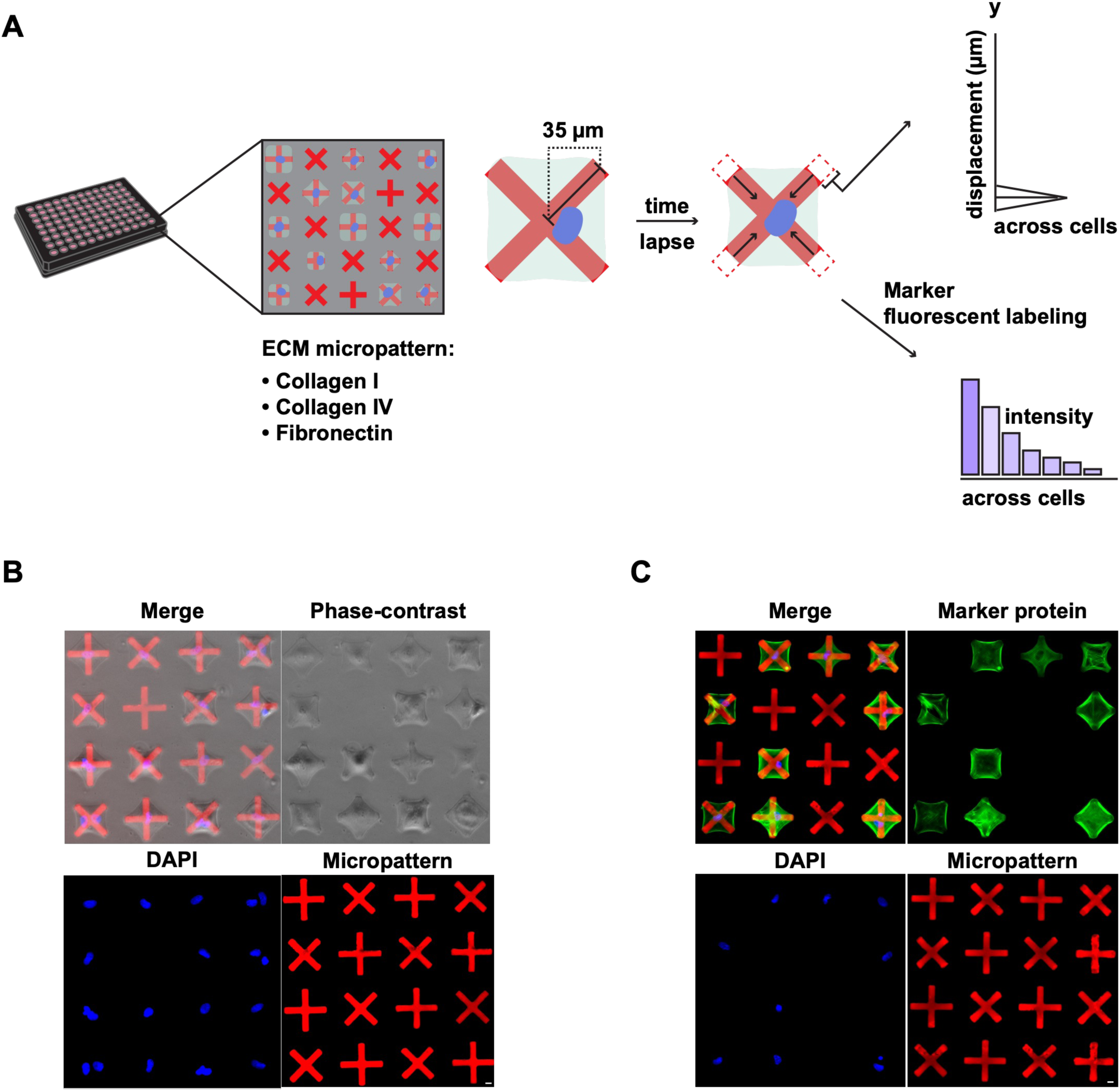
Overview of the single-cell ECM micropattern contractility assay. **(A)** Primary human trabecular meshwork (HTM) cells are seeded onto X-shaped extracellular matrix (ECM) micropatterns composed of collagen I, collagen IV, or fibronectin, with one cell per micropattern. Time-lapse imaging captures micropattern deformation, with inward displacement of the four arms averaged to obtain a single-cell contractile displacement value. Across cells, these measurements form a distribution of contractile behavior. Intracellular marker protein fluorescence staining from the same cells provides a complementary molecular readout. **(B)** Representative HTM cells adhered to ECM micropatterns, shown as merged fluorescence and phase-contrast images and individual channels for nuclear staining (Hoechst) and ECM micropatterns. **(C)** Representative HTM cells following marker protein staining, shown as merged images and individual channels for F-actin (Alexa Fluor 488–phalloidin), nuclear staining (Hoechst), and ECM micropatterns. Scale bar, 10 µm.

## Materials and Methods

### HTM Cell Isolation and Culture

The procurement and use of human eye tissues were approved by the University of California, Los Angeles Institutional Review Board, and all the experiments were performed in accordance with the tenets of the Declaration of Helsinki for the use of human tissues. Primary HTM cells were isolated from healthy corneoscleral rims discarded after transplant surgery as previously described, and cultured according to established protocols^12,13^. Thirteen (13) HTM cell strains (Supplementary Table 1) were used for the experiments in this study between passages 1-3. All HTM donor cell strains were validated by DEX (100 nM; DEX; SIGMA) induced upregulation of myocilin via qRT-PCR. Different combinations of HTM cell strains were used per experiment, with at least three technical replicates provided sufficient material was available. HTM cells were cultured in low-glucose DMEM (1g/L-D glucose; GlutaMAXTM Supplement; 110 mg/L Sodium Pyruvate; Gibco; Thermo Fisher), containing 1X antibiotic-antimycotic (Antibiotic-Antimycotic 100X; Gibco; Thermo Fisher), supplemented with 10% fetal bovine serum (Premium Grade FBS; VWR), and maintained at 37°C in a humidified atmosphere with 5% CO_2_.

### HTM Cell Treatments

Primary human trabecular meshwork (HTM) cells were seeded at a density of 1.5 × 10 cells/cm² onto tissue culture vessels coated with 0.5% gelatin (Type B; Sigma-Aldrich). Upon reaching confluence, cells were serum-starved for 24 h and subsequently treated for 5 days in serum-free medium with vehicle control (0.05% DMSO; Sigma-Aldrich), dexamethasone (DEX; 100 nM; Sigma-Aldrich), the Rho-associated kinase (ROCK) inhibitor Y-27632 (10 µM; Bio-Techne), or combinations thereof. Following treatment, cells were harvested for quantitative RT-PCR, immunocytochemistry, or single-cell contractility analysis.

### Micropattern-based Single-Cell Contractility Assay (FLECS)

Single-cell contractility was quantified using the Fluorescently Labeled Elastomeric Contractible Surfaces (FLECS) assay as previously described^10,11^. Ninety-six-well FLECS plates (Forcyte Biotechnologies) containing red fluorescent, X-shaped ECM micropatterns of defined composition and substrate stiffness were used. Each micropattern measured 70 µm from tip to tip (35 µm per arm from the central junction). To assess ECM-dependent contractility, cells were seeded onto 4–8 kPa substrates micropatterned with collagen I (Cat. No. 96-COL1-AF546-X70-Q1.2), collagen IV (Cat. No. 96-HC4R-X1-QB12), or fibronectin (Cat. No. 96-FN-AF546-X70-Q1.2). To evaluate stiffness-dependent responses, collagen IV micropatterned substrates with stiffnesses of 4–8 kPa (Cat. No. 96-HC4R-X1-QB12), 16–20 kPa (Cat. No. 96-HC4R-X1-QC12), or 40–60 kPa (Cat. No. 96-HC4R-X1-QC16) were used. Unless otherwise indicated, all experiments were performed on collagen IV-coated 4–8 kPa FLECS substrates.

Before cell seeding, phosphate-buffered saline (PBS) was aspirated from each well, and the micropatterns were washed and equilibrated with 100 µL of HTM growth medium containing 10% fetal bovine serum (FBS) at 37°C and 5% CO₂. HTM cells were washed, trypsinized, resuspended in 10% FBS growth medium, passed through a cell strainer, and counted. A total of 3,200 cells were seeded per well in 100 µL of medium supplemented with Hoechst 33342 (1 µg/mL; Thermo Fisher Scientific) for live nuclear staining. Following seeding, plates were incubated for 30 min at room temperature to promote uniform cell settling and then for 30 min at 37°C in a humidified incubator with 5% CO₂ before imaging. Live-cell imaging was performed immediately using an automated ImageXpress Micro XL High-Content Imaging System (Molecular Devices) maintained at 37°C and 5% CO₂. A 4× objective was used to quantify single-cell contractile displacement, whereas a 10× objective was used for imaging immunolabeled cells on ECM micropatterns.

### Image Processing and Analysis

For images acquired for single-cell contractility measurements, raw single-channel TIFF stacks (micropattern and nuclei channels) were processed separately in ImageJ (NIH) to uniformly adjust brightness and contrast for pattern and nucleus visualization. No fluorescence intensity measurements were derived from these images. Single-channel stacks were then analyzed using a proprietary FLECS pipeline provided by Forcyte Biotechnologies and implemented on the Biodock AI platform, a web-based deep AI platform for image analysis. Using standardized file naming, the pipeline automatically paired micropattern and nuclei images from the same field of view, enabling assignment of nuclei to individual micropatterns. Each micropattern consisted of four linear arms extending 35 µm from a central junction. A predefined template was used to detect all patterns in each frame, and for each pattern the distances from the center to the tip of all four arms were measured and averaged to yield a single data point. Patterns were classified as single-cell, multinucleated, or multi-cell based on the presence of stained nuclei. Cell-free patterns were used as controls of pattern dimensions, and only single-cell patterns were retained for downstream analysis. For each experimental condition, pattern dimensions were zeroed to the median value of the corresponding control patterns to yield the net inward displacement, such that the single-cell contraction measurement represents the change in micropattern dimensions (net reduction in arm length) relative to the undisturbed control patterns.

For the contraction versus marker protein fluorescence intensity studies, images were processed using a custom analysis pipeline in Biodock AI (Version 13). Three-channel composite images (ECM micropattern: red; marker: green; nuclei: blue) were exported as single overlaid TIFF files. An AI-based segmentation model, trained on manually annotated images, identified three object classes: nuclei, micropatterns, and square box regions surrounding each pattern. Large fields of view were tiled at multiple scales (smaller tiles for nuclei, larger tiles for patterns and box regions), and only tiles annotated as fully labeled were used for training. A hierarchical assignment linked nuclei to micropatterns and micropatterns to box regions using a predefined overlap threshold, enabling calculation of the number of nuclei per pattern within each box. Marker fluorescence intensity was quantified within the box regions, which were centered on each micropattern and sized to encompass the associated cell area; intensity metrics from each box were then paired with the contraction measurement from the enclosed micropattern. Marker positivity was assessed using raw green-channel fluorescence intensity measured within AI-segmented objects. Background fluorescence was established using no-primary-antibody control images, and objects with green-channel intensity at or below the control-derived background threshold were classified as marker-negative. To avoid potential confounding effects, all correlation analyses between marker abundance and contractility were restricted to marker-positive cells (Supplementary Figure 4, Supplementary Table 2).

For each segmented object, the pipeline computed geometric features (e.g. area, major-axis length, curved length) and per-channel intensity metrics. All quantitative features were extracted from the original raw image files (brightness and contrast were adjusted only in visualization copies), and results were exported as CSV files for downstream data processing and statistical analysis in R.

### Immunofluorescent Staining on FLECS Plates

HTM cells were rinsed with PBS and fixed in pre-warmed 3.7% paraformaldehyde for 20 min. Cells were washed with PBS and permeabilized with 0.25% Triton X-100 for 10 min at room temperature with nutation. They were then washed with PBS and blocked with 5% goat serum for 30 min at room temperature with nutation. HTM cells were incubated overnight at 4°C in antibody dilution buffer (1% BSA, 1% goat serum, 0.3% Triton X-100 in PBS) with nutation in the following primary antibodies: rabbit anti-myocilin (1:500; Stamer et al., 1995), mouse anti-ANGPTL7 (1:200; AdipoGen Life Sciences; cat# KAIROS 108-4), mouse anti–α-SMA (1:500; Millipore Sigma; cat# A5228), rabbit anti–phospho-MLC2 (1:200; Cell Signaling Technology; cat# 3674), rabbit anti-fibronectin (1:500; Abcam; cat# ab2413), rabbit anti-FABP4 (1:100; ABclonal; cat# A11481), mouse anti-TMEFF2 (1:100; Santa Cruz; cat# sc-376175), rabbit anti-CHI3L1 (1:100; ABclonal; cat# A11510), and mouse anti-αB crystallin (1:100; StressMarq; cat# SMC-159). Alexa Fluor 488 goat anti-rabbit or goat anti-mouse IgG (H+L), cross-adsorbed secondary antibodies (1:500; Invitrogen), were used for signal detection, and Hoechst 33342 was used for nuclear staining (1 µg/mL; Thermo Fisher). F-actin was stained with Alexa Fluor 488 phalloidin (1:400; Invitrogen) on the day of secondary antibody incubation.

### Quantitative Reverse Transcription PCR

Total RNA was isolated from cells using the RNeasy kit (Qiagen, Valencia, CA) according to the manufacturer’s instructions. RNA concentration was measured using a Nanodrop spectrophotometer (Thermo Fisher, Waltham, MA). cDNA was synthesized from total RNA using the QuantiTect Reverse Transcription kit (Qiagen, Germantown, MD) following the manufacturer’s instructions. Quantitative PCR was performed on a realplex² thermocycler (Eppendorf, Hauppauge, NY) using TaqMan assays for human GAPDH (Hs02758991_g1) and human MYOC (Hs00165345_m1). Reactions were run in technical triplicates. Data were normalized to GAPDH and analyzed using the ΔΔCt method.

### Statistical Analysis via Non-Parametric Resampling

All statistical analyses and visualizations were performed using the R statistical computing environment (v4.4.3). Due to the pronounced skew and non-normality of the single-cell contractility data, nonparametric resampling methods were used for all inferential statistics. This approach permitted the use of the median rather than the mean as a robust measure of central tendency while explicitly accounting for the hierarchical structure of the data (cells nested within wells, nested within human donor cell strains). Statistical significance was evaluated at an α level of 0.05, unless otherwise indicated.

To capture the heterogeneity and skew of single-cell populations, summary statistics are reported as medians and interquartile ranges (IQR). Confidence intervals were estimated using bootstrap resampling appropriate to the experimental design. Cluster bootstrap resampling (donors) was applied to cell-level analyses, nested bootstrap resampling (donors followed by wells) was used for pooled correlation analyses, and bootstrap resampling of wells within donors was used for donor-specific correlation analyses. In all cases, 95% confidence intervals were obtained from the empirical percentiles of the bootstrap distribution across 10,000 iterations. Hypothesis testing was performed using permutation tests without assuming a specific parametric distribution. Labels were permuted without replacement while preserving the experimental design hierarchy. Test statistics were recomputed for 10,000 random permutations to generate empirical null distributions, from which one– or two-sided p-values were calculated as appropriate. Multiple comparisons were adjusted using either the Holm–Bonferroni or Benjamini–Hochberg procedure, as specified for each analysis. Associations between single-cell contractile displacement and marker fluorescence intensity were quantified using Spearman’s rank correlation coefficient (ρ). Correlation analyses are reported as effect estimates with 95% confidence intervals rather than hypothesis tests.

Instead of a classical one-way ANOVA, which assumes normally distributed data and uses group means to compute the F statistic, a nonparametric permutation-based omnibus test was used. Between-group variability of group medians around the grand median was contrasted with within-group variability of individual observations around their respective group medians. Statistical significance was assessed from the empirical permutation distribution generated for 10,000 random permutations.

When an omnibus permutation test indicated a significant global effect, post hoc pairwise permutation tests were performed. For repeated time-resolved comparisons, a Max-T correction was applied using the maximum test statistic across all time intervals from each permutation to construct the reference null distribution, thereby controlling the familywise error rate.

## Results

### ECM identity organizes single-cell contractile dynamics

To determine how extracellular matrix (ECM) identity shapes human trabecular meshwork (HTM) cell contractile behavior, we quantified time-resolved single-cell contractile displacement on collagen IV, collagen I, or fibronectin micropatterns approximating normal stiffness (4–8 kPa)^14,15^. We found that ECM identity influenced both the magnitude and temporal organization of single-cell contractile behavior (Fig. 2).

**Figure 2.**
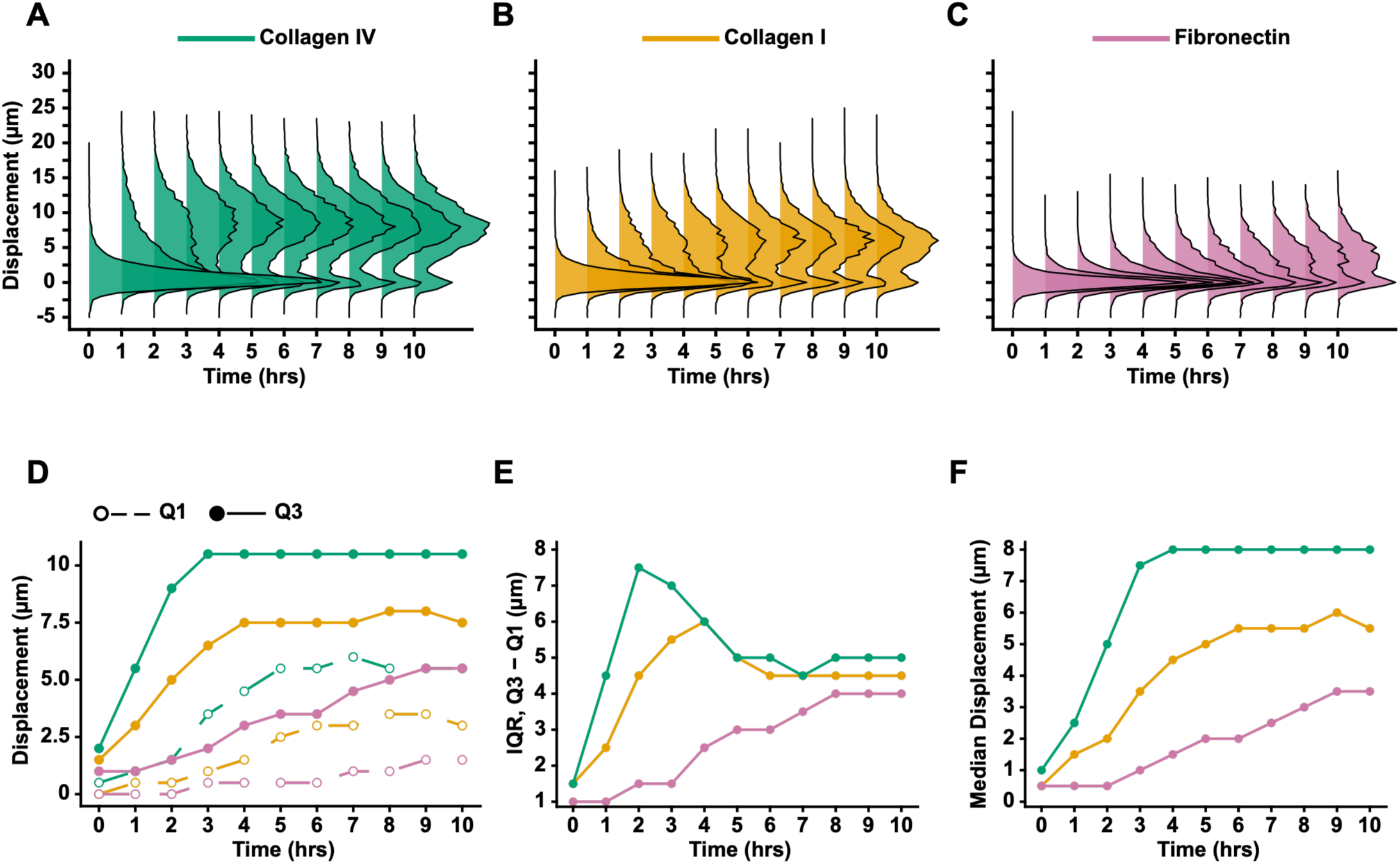
Extracellular matrix identity shapes the temporal organization of single-cell HTM contractility. Primary human trabecular meshwork (HTM) cells were seeded on collagen IV, collagen I, or fibronectin micropatterns, and single-cell contractile displacement was measured over 10 h. **(A–C)** Single-cell contractile displacement distributions on collagen IV (**A**, green), collagen I (**B**, yellow), and fibronectin (**C**, magenta) over 10 h. **(D)** First (Q1; 25th percentile) and third (Q3; 75th percentile) quartiles of contractile displacement over time. Dashed lines with open circles indicate Q1; solid lines with closed circles indicate Q3. **(E)** Interquartile range (IQR; Q3 − Q1) of contractile displacement over time. **(F)** Median contractile displacement over time. Data are from one human donor strain (TM408P1), with a minimum of 3,000 cells analyzed per timepoint per ECM condition.

HTM cells adherent to collagen IV showed rapid distributional expansion followed by convergence toward higher contractile displacement (Fig. 2A). By 2 h, the distribution had expanded substantially, with a first quartile (Q1; 25th percentile) of 1.5 µm and a third quartile (Q3; 75th percentile) of 9 µm (Fig. 2D). This corresponded to an interquartile range (IQR; Q3 − Q1) of 7.5 µm, the largest observed over the 10-h period (Fig. 2E). Subsequently, the distribution narrowed as Q3 stabilized at 10.5 µm, while Q1 continued to increase, reaching 5.5 µm by 10 h (Fig. 2D,E). Median contractile displacement reached 8 µm by 4 h, the highest observed across the three ECM conditions (Fig. 2F). Thus, as the upper quartile stabilized and the lower quartile shifted toward higher displacement, the distribution converged toward a high-displacement, increasingly symmetric state.

HTM cells adherent to collagen I showed a slower but sustained contractile trajectory, with lower displacement and less pronounced distributional convergence compared with collagen IV (Fig. 2B). The distribution expanded through 4 h, reaching an IQR of 6 µm, before narrowing to 4.5 µm by 6 h (Fig. 2E). This narrowing reflected stabilization of Q3 near 7.5 µm, approximately 3 µm below collagen IV, while Q1 continued to increase, reaching 3 µm by 10 h (Fig. 2D). Median contractile displacement reached 5.5 µm at 10 h, remaining lower than on collagen IV (Fig. 2F). Thus, collagen I displayed slower distributional expansion and less pronounced convergence toward higher displacement than collagen IV.

In contrast, HTM cells adherent to fibronectin maintained comparatively low displacement and showed no early expansion of the distribution, which instead broadened progressively over time (Fig. 2C). Median displacement increased from 0.5 µm at baseline to 3.5 µm at 10 h (Fig. 2F), while the IQR widened progressively from 1 µm to 4 µm over the same period (Fig. 2E). This increasing dispersion reflected greater extension of the upper than lower quartile of the distribution (Fig. 2D). Thus, fibronectin produced a fundamentally distinct, lower-magnitude and progressively broadening distributional response rather than a temporally delayed version of the collagen-like expansion and convergence.

Collectively, our findings show that ECM identity shapes not only the magnitude of single-cell HTM contractility but, importantly, the temporal organization of the response. On both collagen types, rapid distributional expansion gave way to convergence toward higher displacement, whereas on fibronectin, lower displacement was accompanied by progressive broadening throughout the observation period. Together, these results demonstrate that ECM identity organizes single-cell contractile behavior into distinct temporal and distributional patterns rather than simply scaling the magnitude or kinetics of a common mechanical response.

### Increasing ECM stiffness restricts single-cell contractile behavior

Having established that ECM identity shapes the temporal organization of single-cell HTM contractile behavior, we next investigated how the magnitude and distribution of single-cell contractile displacement change with increasing substrate stiffness. To model the progressive stiffening of the trabecular meshwork that accompanies glaucoma progression, HTM cells pretreated with vehicle or 100 nM DEX were seeded on collagen IV micropatterns spanning normal (4–8 kPa), intermediate (16–20 kPa), and pathological (40–60 kPa) stiffnesses. Single-cell contractile displacement was measured over 10 h.

Increasing substrate stiffness progressively constrained the range and organization of single-cell HTM contractile behavior (Fig. 3). Under normal stiffness, vehicle– and DEX-treated cells displayed broad displacement distributions spanning minimally displaced cells to high-displacement behavior exceeding 10 µm at later time points. Within this heterogeneous population, DEX-treated cells formed a pronounced higher-displacement shoulder relative to vehicle, consistent with enhanced contractile output within a portion of the population rather than a uniform population-wide shift (Fig. 3A, D, G). Increasing stiffness progressively reshaped this response. At intermediate stiffness, the overall displacement range narrowed, and high-displacement behavior became less prominent, although DEX-treated cells retained a modest extension of the upper range relative to vehicle (Fig. 3B, E, H). At pathological stiffness, this restriction became more pronounced, with both treatment groups concentrated toward lower displacement and little separation between vehicle and DEX. Notably, pathological stiffness did not eliminate contractile behavior, as a residual range of displacement remained evident despite the marked loss of high-displacement behavior (Fig. 3C, F, I). Together, these distributions show that increasing stiffness progressively restricts the range of single-cell contractile behavior while attenuating the heterogeneous response to DEX.

**Figure 3.**
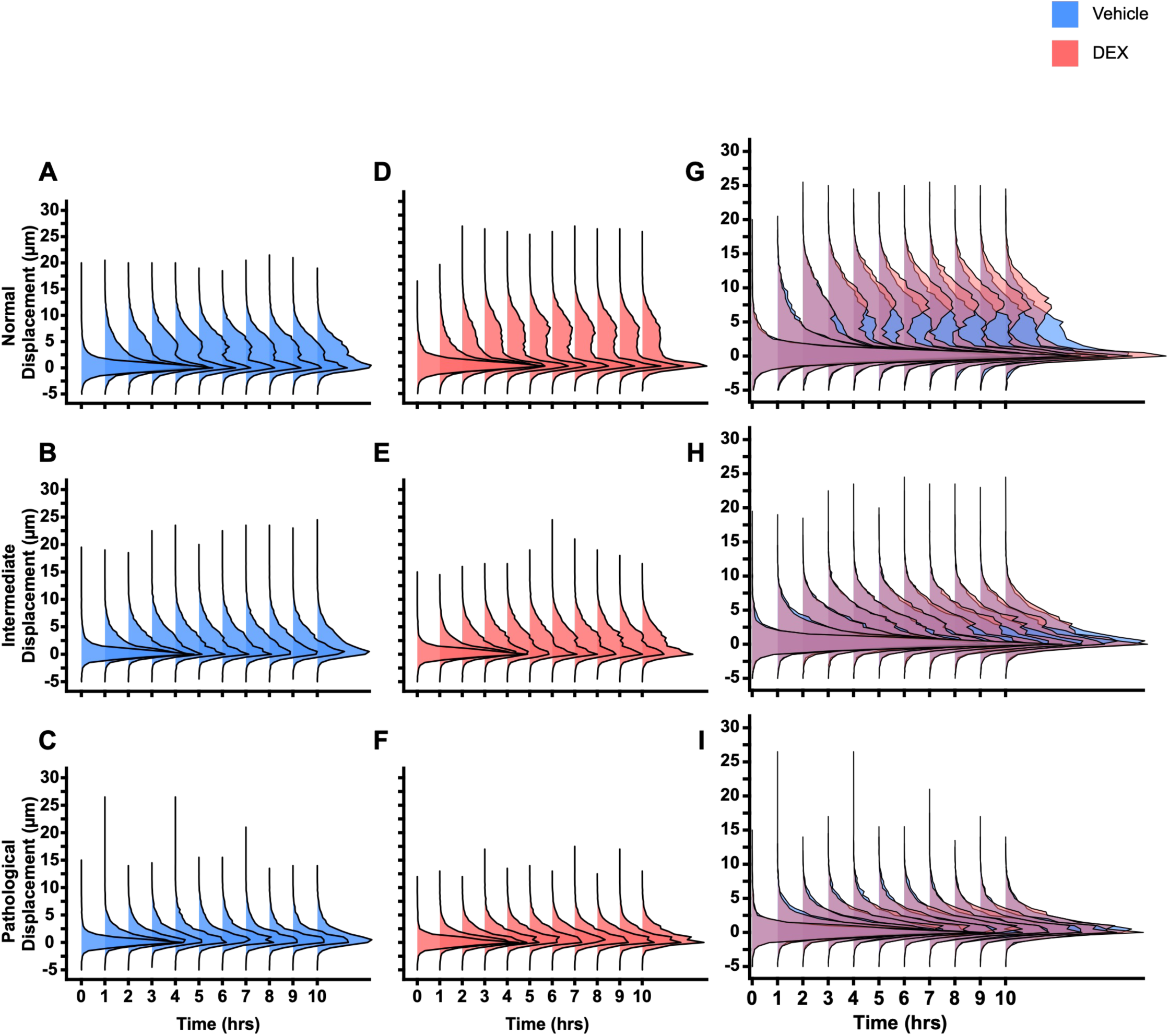
Substrate stiffness progressively constrains single-cell contractile heterogeneity. Single-cell contractile displacement distributions over time on collagen IV micropatterned substrates of normal (4–8 kPa), intermediate (16–20 kPa), and pathological (40–60 kPa) stiffness. Cells were pre-treated for 5 days with vehicle (0.05% DMSO) or 100 nM DEX. Columns show vehicle-treated **(A–C)**, DEX-treated **(D–F)**, and overlaid vehicle and DEX **(G–I)** distributions. Rows show normal **(A, D, G)**, intermediate **(B, E, H)**, and pathological **(C, F, I)** stiffness conditions. Each ridge plot shows single-cell contractile displacement distributions at hourly timepoints from 0 to 10 h. Data represent N = 4 human donors (TM306, TM318, TM337, TM340) across all stiffness conditions, with a minimum of 10,000 cells analyzed per timepoint per condition.

This stiffness-dependent restriction of single-cell behavior was accompanied by a progressive reduction in contractile displacement across donor strains (Fig. 4A–C). On normal stiffness, median displacement increased over the first several hours before plateauing at 4.75 µm in vehicle-treated cells and 6.25 µm in DEX-treated cells by 5 h (Fig. 4A). At intermediate stiffness, the magnitude of contraction was substantially reduced, with vehicle– and DEX-treated cells reaching lower plateaus of 2.0 and 2.5 µm, respectively, by 4 h (Fig. 4B). This reduction was most pronounced on pathological stiffness, where both treatment groups reached median displacements of only 1.25 µm by 3 h (Fig. 4C). Thus, the progressive restriction of the single-cell displacement range observed across the full distributions was accompanied by a corresponding reduction in the magnitude of contractile displacement across donor strains.

**Figure 4.**
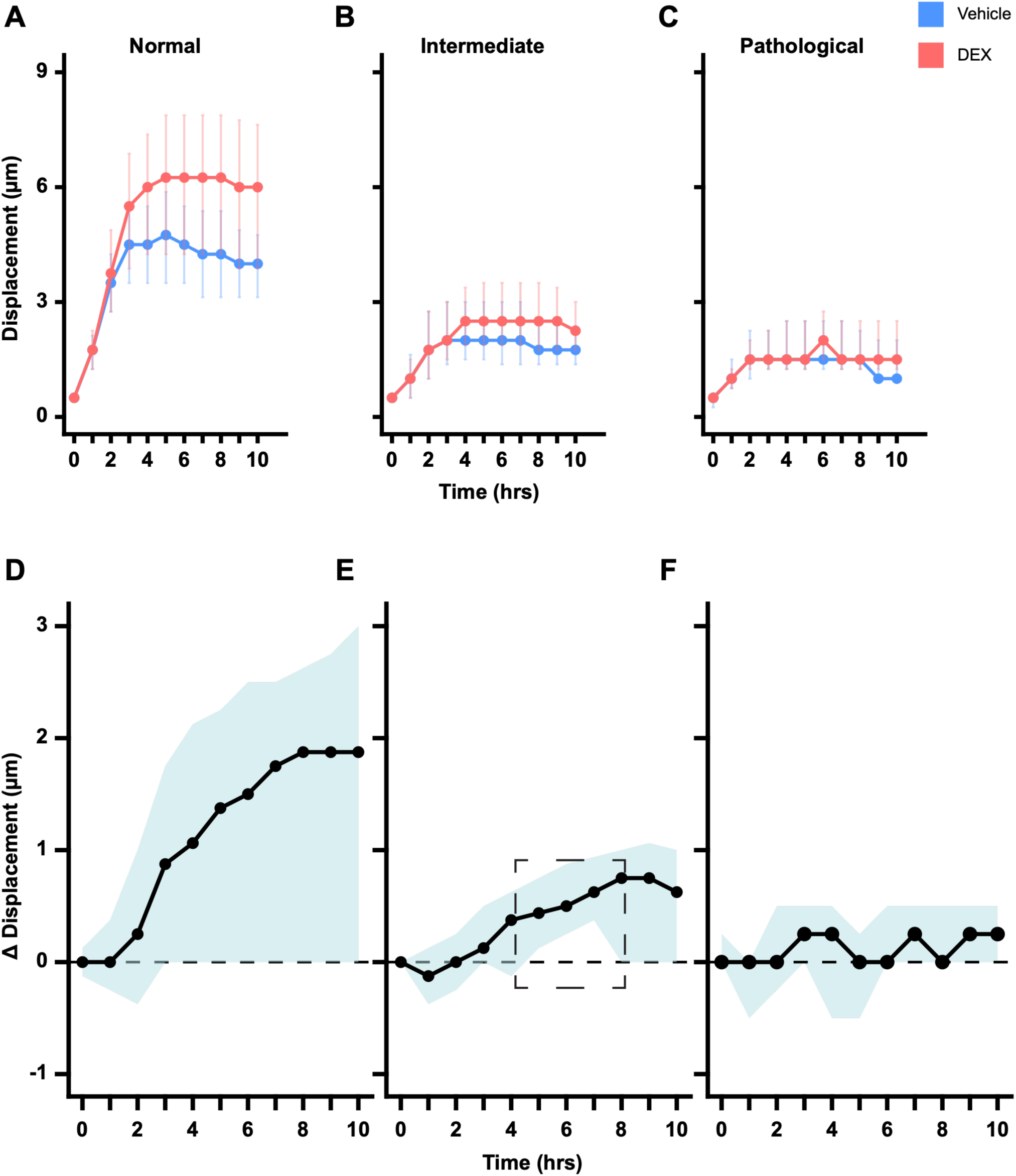
Substrate stiffness constrains single-cell contractile displacement and DEX responsiveness in HTM cells. HTM cells pre-treated for 5 days with vehicle (0.05% DMSO) or 100 nM DEX were seeded on collagen IV micropatterned substrates of normal (4–8 kPa), intermediate (16–20 kPa), or pathological (40–60 kPa) stiffness. Columns show normal **(A, D)**, intermediate **(B, E)**, and pathological **(C, F)** stiffness conditions. **(A–C)** Median contractile displacement trajectories for vehicle and DEX treatment, with IQR. **(D–F)** Time-resolved differences in median contractile displacement between DEX and vehicle [Δ(DEX − VEH)], with shaded 95% cluster-bootstrap confidence intervals (CIs). The dashed box in E indicates timepoints at which the 95% CI excluded zero. **(A–C)** N = 4 human donors (TM306, TM318, TM337, TM340) for all stiffness conditions. **(D–F)** N = 4 donors for normal **(D)** and intermediate **(E)** stiffness and N = 3 donors (TM306, TM318, TM337) for pathological stiffness (**F**; see Methods). A minimum of 10,000 cells were analyzed per timepoint per condition.

The DEX-associated effect on contractile displacement was also progressively attenuated with increasing substrate stiffness (Fig. 4D–F). On normal stiffness, the DEX-associated difference in median contractile displacement within donor strains [Δ(DEX−VEH)] increased over time, reaching 1.875 µm at 8 h, although the 95% cluster-bootstrap confidence interval (CI) included zero from 3 h onward, reflecting substantial variation among donor strains (Fig. 4D). At intermediate stiffness, the DEX-associated difference was smaller, reaching 0.75 µm at 8 h, with 95% CIs excluding zero from 5–7 h (Fig. 4E). On pathological stiffness, the difference between treatment groups remained minimal throughout the observation period, never exceeding 0.25 µm, with 95% CIs overlapping zero at all timepoints (Fig. 4F). Together, these results show that the enhanced contractile displacement associated with DEX is progressively attenuated as substrate stiffness increases.

### DEX sustains contractile activity on normal stiffness

To determine whether DEX alters the kinetics of contractile displacement, we quantified hourly displacement rates across donor strains using the same cohort analyzed in Figs. 3 and 4. On normal stiffness, DEX maintained a higher displacement rate than vehicle, whereas no significant treatment differences were detected on intermediate or pathological stiffness (Fig. 5A–C).

**Figure 5.**
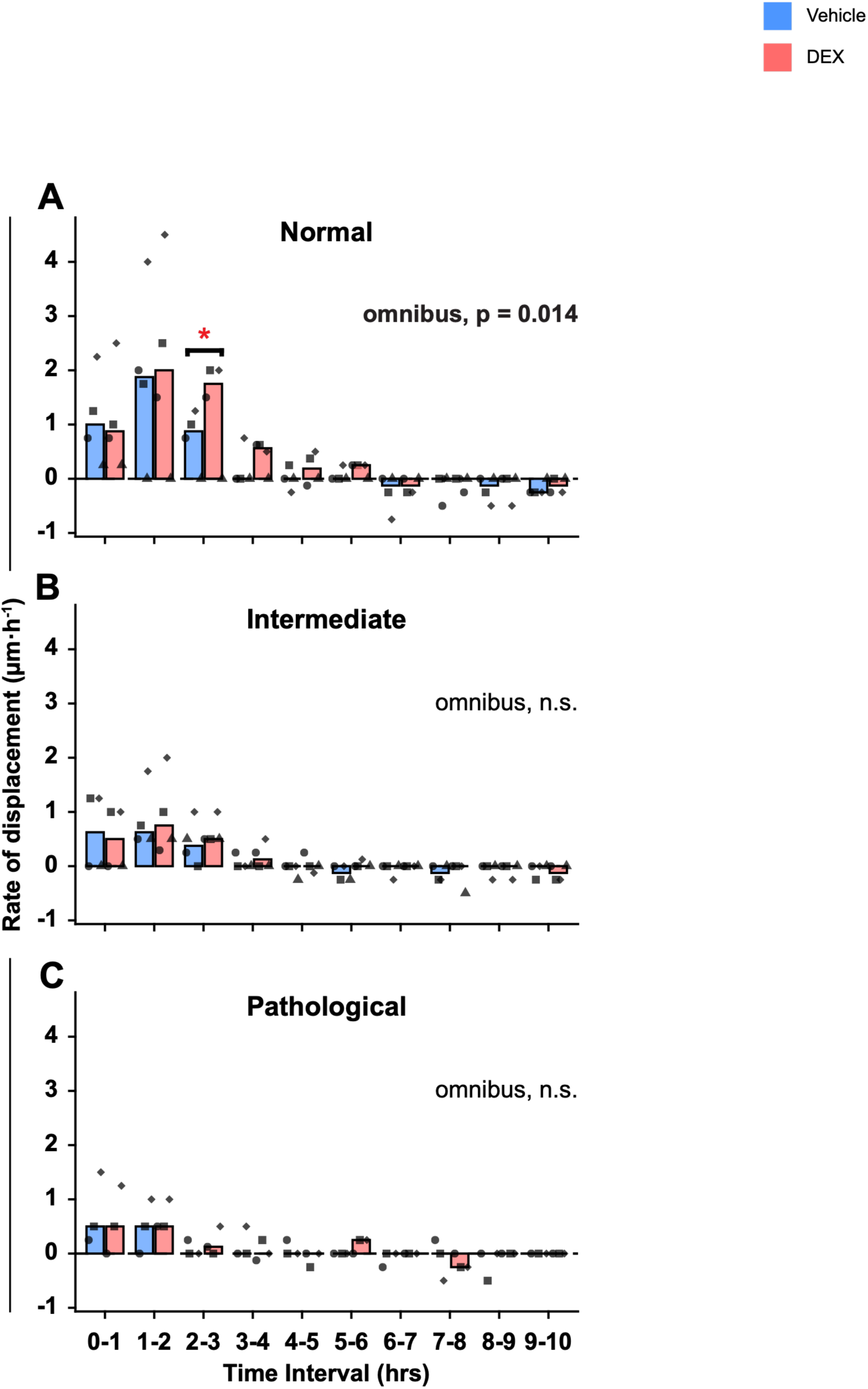
Substrate stiffness gates DEX-dependent contractile dynamics in HTM cells. HTM cells pre-treated for 5 days with vehicle (0.05% DMSO) or 100 nM DEX were seeded on collagen IV micropatterned substrates of normal (4–8 kPa), intermediate (16–20 kPa), or pathological (40–60 kPa) stiffness. **(A–C)** Median rate of displacement (µm h⁻¹) across donor strains for each 1-h interval, calculated from the interval-by-interval change in well-level median cell displacement; symbols indicate individual donor strains. A one-sided omnibus permutation test (DEX > vehicle) detected a treatment effect on normal stiffness (P = 0.014; Holm–Bonferroni corrected). Post hoc pairwise permutation testing with Max-T correction identified a DEX-associated difference at the 2–3 h interval on normal stiffness (P = 0.029). No significant differences were detected on intermediate or pathological stiffness. N = 4 human donor strains (TM306, TM318, TM337, TM340) for normal **(A)** and intermediate **(B)** stiffness and N = 3 human donor strains (TM306, TM318, TM337) for pathological stiffness (**C**; see Methods), with a minimum of 10,000 cells analyzed per timepoint per condition.

On normal stiffness, vehicle– and DEX-treated cells showed similar early kinetics, reaching peak displacement rates of 1.87 and 2.0 µm·h⁻¹, respectively, between 1 and 2 h (Fig. 5A). Their trajectories diverged after this shared peak as vehicle-treated cells more readily decelerated, whereas displacement rate remained elevated under DEX. Between 2 and 3 h, the median displacement rate declined to 0.87 µm·h⁻¹ under vehicle but remained at 1.75 µm·h⁻¹ under DEX, representing the interval of greatest separation between treatment groups (permutation test, p = 0.029; Max-T corrected). Thereafter, displacement rates declined under both conditions.

Increasing substrate stiffness progressively reduced the magnitude of the kinetic response and diminished the separation between vehicle and DEX. On intermediate stiffness, peak displacement rates were substantially lower than on normal stiffness, reaching 0.62 and 0.75 µm·h⁻¹ under vehicle and DEX, respectively, before declining over subsequent intervals with only modest separation between treatment groups (Fig. 5B). On pathological stiffness, displacement rates remained low under both conditions, reaching a peak of only 0.5 µm·h⁻¹ under both vehicle and DEX, with little separation between their subsequent trajectories (Fig. 5C). Thus, rather than substantially increasing the initial peak displacement rate, DEX prolonged elevated contractile activity on normal stiffness, an effect that was progressively constrained as stiffness increased.

We next examined whether DEX-associated displacement responses persisted into the relaxation phase using an independent cohort of three HTM cell strains monitored from 10 to 16 h on normal-stiffness substrates (Fig. 6). Although displacement declined under both conditions, DEX-treated populations consistently retained a more extended high-displacement range. For TM267 and TM318 cells, DEX-treated distributions remained shifted toward higher displacement relative to vehicle, consistent with their elevated median trajectories (Fig. 6A–D). In contrast, median displacement of TM271 cells fell below vehicle by 16 h, while the DEX-treated population continued to exhibit a high-displacement tail (Fig. 6E,F). Notably, the upper quartile (Q3) remained higher under DEX across all three strains at every time point, indicating persistent high-displacement behavior despite donor-specific differences in median responses.

**Figure 6.**
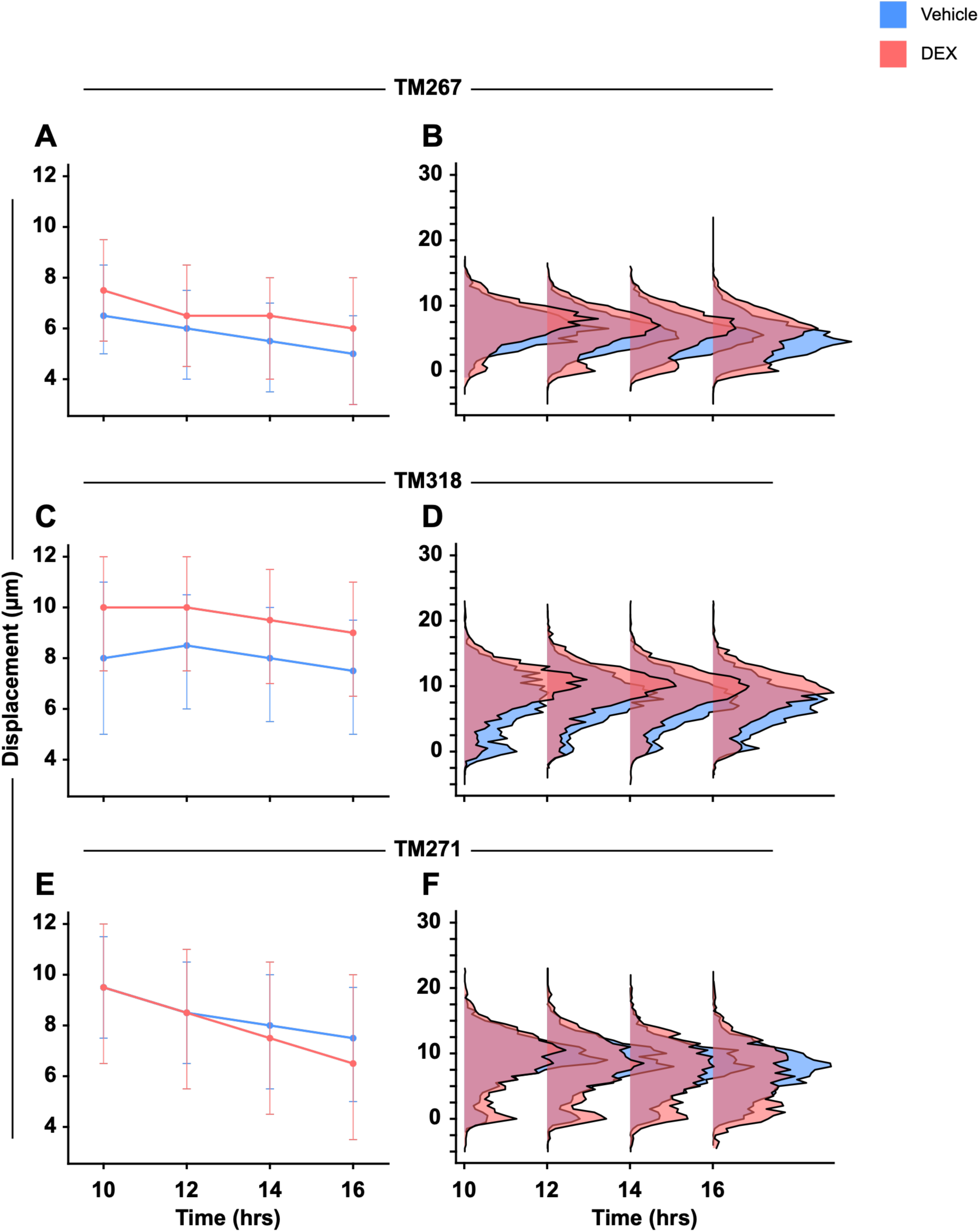
Median contractile displacement with interquartile range on normal-stiffness substrates for three independent TM strains. Corresponding overlaid single-cell displacement distributions for vehicle and DEX at each timepoint (IQR at 10, 12, 14, and 16 h). Three human donor TM strains (TM267, TM318, TM271), with a minimum of 6,500 cells analyzed per timepoint per condition.

An additional HTM cell strain (TM317) exhibited a persistent bimodal displacement distribution that could not be adequately characterized by a single measure of central tendency and was therefore analyzed separately (Supplementary Fig. 2). Despite this distinct distributional profile, DEX similarly maintained an extended high-displacement range throughout the relaxation phase.

Together, these findings demonstrate that DEX-associated high-displacement behavior persists into the relaxation phase on normal-stiffness substrates (4–8 kPa), despite substantial donor-specific heterogeneity. Combined with our observations across substrate stiffnesses, these results further suggest that increasing stiffness progressively attenuates the observable DEX-associated displacement response. Importantly, the prolonged response is most evident in the high-displacement portion of the population rather than as a uniform increase in displacement across all HTM cells.

## ROCK inhibition restricts contractile magnitude and heterogeneity

Topical Rho-associated kinase (ROCK) inhibitors are increasingly used to lower intraocular pressure in patients with ocular hypertension and glaucoma16^16,17^. Because ROCK signaling regulates actomyosin tension and aqueous humor outflow, we examined how ROCK inhibition alters single-cell contractility with and without DEX. To do this, we pretreated HTM cells from four donor-derived strains with vehicle, 100 nM DEX, 10 µM Y-27632 (ROCK inhibitor; ROCKi), or DEX plus Y-27632 (DEX + ROCKi), and seeded them on collagen IV micropatterns of normal stiffness (4–8 kPa) for a 10-h observation period (Fig. 7).

**Figure 7.**
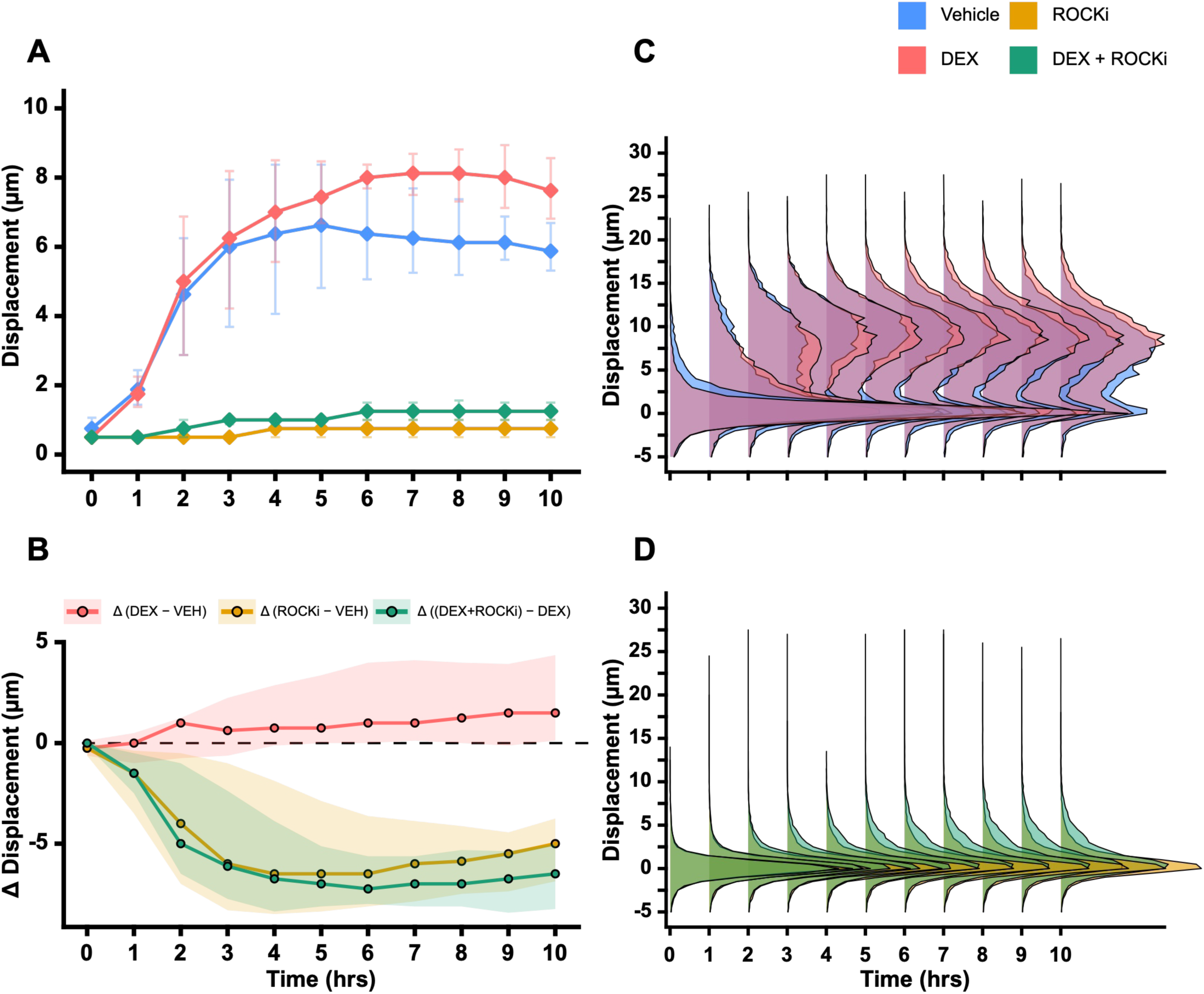
ROCK inhibition constrains basal and DEX-enhanced single-cell contractile behavior. HTM cells were pre-treated for 5 days with vehicle (0.05% DMSO), 100 nM dexamethasone (DEX), 10 µM ROCK 1/2 inhibitor (Y-27632; ROCKi), or DEX and ROCKi (DEX+ROCKi), then seeded on collagen IV micropatterns of normal stiffness (4–8 kPa). Single-cell contractile displacement was measured over 10 h. **(A)** Median contractile displacement trajectories for each treatment condition, with error bars indicating inter-donor IQR. **(B)** Time-resolved differences in median contractile displacement [Δ(DEX − VEH), Δ(ROCKi − VEH), and Δ((DEX+ROCKi) − DEX)], with shaded 95% confidence intervals (CIs). **(C)** Overlaid single-cell displacement distributions for vehicle– and DEX-treated cells. **(D)** Overlaid single-cell displacement distributions for ROCKi– and DEX+ROCKi-treated cells. Data represent N = 4 human donor strains (TM349, TM402, TM403, TM405), with a minimum of 14,000 cells analyzed per timepoint per condition.

ROCK inhibition substantially restricted the range of single-cell contractile displacement. Vehicle– and DEX-treated cells developed broad displacement distributions extending beyond 15 µm, with DEX producing a more extended high-displacement range (Fig. 7C). In contrast, ROCKi-treated cells remained narrowly concentrated at low displacement throughout the observation period (Fig. 7D). DEX + ROCKi produced a similarly restricted distribution, although a modest upper tail emerged at later timepoints (Fig. 7D). Thus, ROCK inhibition fundamentally reorganized the distribution of single-cell contractile displacement, shifting broadly heterogeneous populations toward a restricted, low-displacement range under both basal and DEX-treated conditions.

Consistent with this distributional restriction, ROCK inhibition substantially reduced median contractile displacement across donor strains. Vehicle– and DEX-treated cells showed pronounced time-dependent increases in displacement, reaching medians of approximately 6.75 and 8.25 µm, respectively. In contrast, trajectories remained low and comparatively flat under ROCK inhibition, plateauing at 0.75 µm with ROCKi and reaching only 1.25 µm with DEX + ROCKi by 10 h (Fig. 7A). Thus, despite a modest late increase with DEX + ROCKi co-treatment, contractile displacement remained substantially lower than under vehicle or DEX alone.

We next quantified these treatment effects across donor strains over time. DEX increased contractile displacement relative to vehicle, with considerable inter-donor variability in response magnitude, consistent with our preceding findings. In contrast, ROCKi diverged from vehicle within the first hour, with 95% cluster-bootstrap CIs excluding zero from 1 h onward and reaching a maximum difference of −6.5 µm between 4 and 6h. A similarly large difference was observed between DEX + ROCKi and DEX, reaching −7.25 µm at 6 h (95% CI: −8.0 to −5.625 µm) and remaining at −6.5 µm at 10 h (Fig. 7B). Together, these comparisons demonstrate that ROCK inhibition produced a rapid and sustained suppression of contractile displacement across donor strains that persisted even in the presence of DEX.

Collectively, these findings demonstrate that ROCK inhibition rapidly and persistently suppresses basal and DEX-associated contractile dynamics. Rather than simply reversing the DEX-associated increase in displacement, ROCK inhibition reorganized the broad heterogeneous distributions observed under both conditions into a distinctly narrow, low-displacement range.

### Contractile heterogeneity is not captured by individual marker abundance

We next asked whether differences in intracellular protein abundance were associated with the heterogeneity in single-cell contractile displacement by simultaneously measuring both features in individual HTM cells. Six markers representing two complementary biological categories were examined: cytoskeletal and ECM-associated proteins implicated in contractile activity, including filamentous actin (F-actin), phosphorylated myosin light chain-2 (p-MLC2), α-smooth muscle actin (α-SMA), and fibronectin (FN); and two glaucoma-associated steroid-responsive biomarkers, myocilin (MYOC) and angiopoietin-like 7 (ANGPTL7), whose relationship to cellular mechanics remains poorly defined (Supplementary Fig. 3).

Consistent with the preceding experiments, DEX treatment significantly increased contractile displacement across all six marker-positive populations (Fig. 8, top marginal). Median treatment effects [Δ(DEX−VEH)] across donors ranged from 0.76 µm in MYOC-positive cells to 3.03 µm in F-actin-positive cells. Across all six markers, DEX also broadened the range of contractile displacement, with high-displacement behavior extending beyond that observed under vehicle.

**Figure 8.**
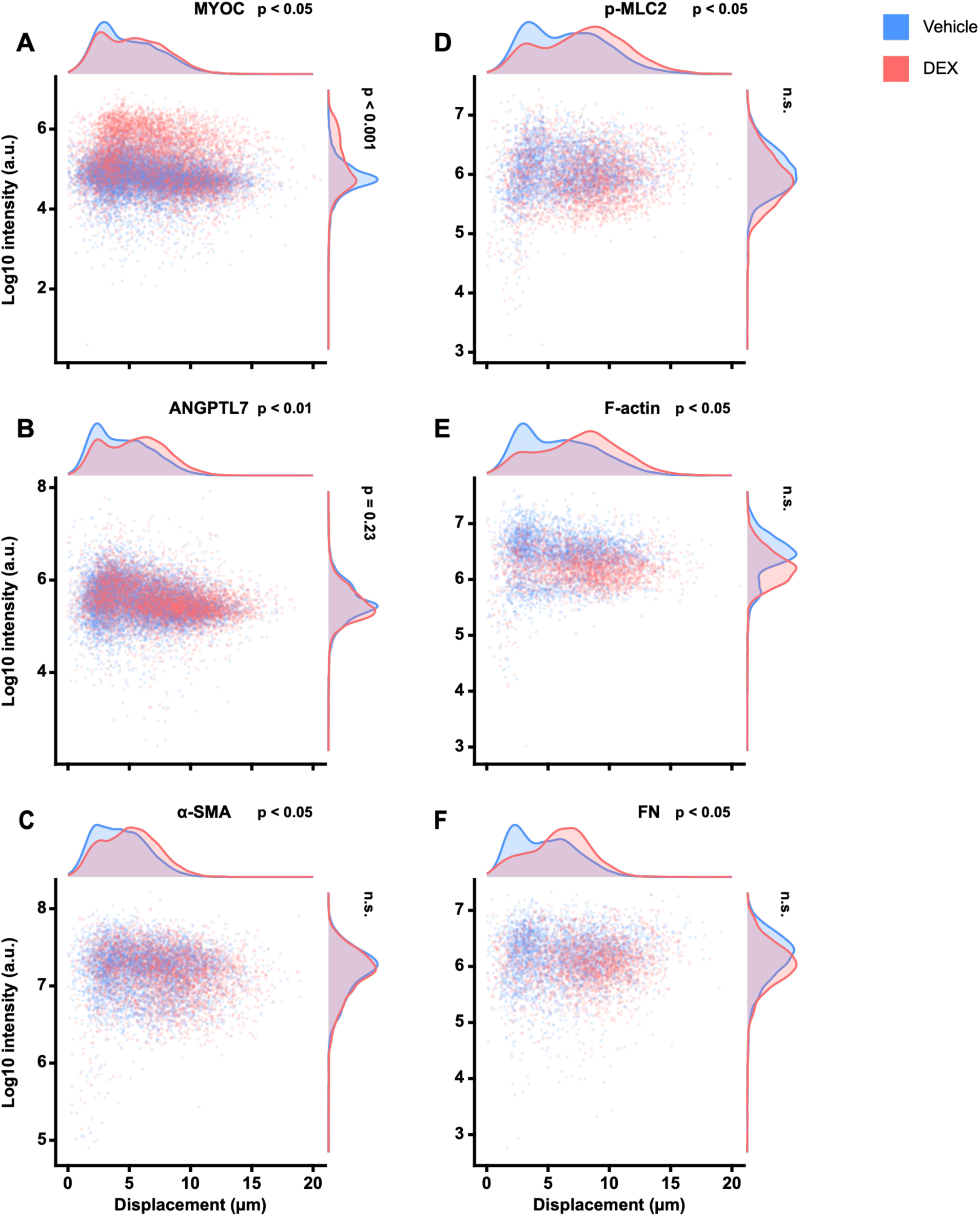
DEX enhances contractile displacement across molecularly heterogeneous HTM cells. Bivariate distributions of single-cell contractile displacement and log10-transformed fluorescence intensity for six protein markers in HTM cells treated with vehicle (0.05% DMSO) or 100 nM DEX. Cells were seeded for 6 h on collagen IV micropatterns of normal stiffness (4–8 kPa) before immunocytochemistry. Each panel shows the bivariate distribution of single-cell measurements with marginal density distributions for contractile displacement (top) and fluorescence intensity (right); y-axis ranges differ among markers. **(A)** myocilin (MYOC), **(B)** angiopoietin-like 7 (ANGPTL7), **(C)** alpha-smooth muscle actin (α-SMA), **(D)** phosphorylated myosin light chain 2 (p-MLC2), **(E)** F-actin, and **(F)** fibronectin (FN). Permutation-derived P values indicate treatment effects on median contractile displacement (top) and mean log10 fluorescence intensity (right). Tests were one-sided and accounted for the hierarchical structure of the data (cells nested within wells nested within donors), with Benjamini–Hochberg false discovery rate correction for multiple testing. N = 4 human donor strains (TM349, TM403, TM404, TM405) for MYOC, ANGPTL7, and α-SMA and N = 3 human donor strains (TM403, TM404, TM405) for p-MLC2, F-actin, and FN, with a minimum of 2,000 cells analyzed per marker per condition.

In contrast to the mechanical response, DEX-associated changes in intracellular protein abundance were marker-dependent (Fig. 8, right marginal). MYOC was the only marker significantly increased by DEX treatment, with a 2.31-fold increase relative to vehicle (BH-adjusted *p < 0.001*). ANGPTL7 exhibited a more modest 1.33-fold increase that did not remain significant after multiple-testing correction, and no significant changes in protein abundance were detected for α-SMA, p-MLC2, F-actin, or FN. Examining both measurements together at the single-cell level revealed minimal correspondence between protein abundance and contractile displacement, with no apparent trend across any of the six markers (Fig. 8, central scatter). Cells with similar protein abundance levels spanned a broad range of contractile displacement, a pattern that persisted across both treatment conditions.

To quantify the relationship between intracellular protein abundance and contractile displacement, we calculated Spearman correlation coefficients (ρ; rho) from paired single-cell measurements using a block-aware framework that preserved the hierarchical structure of the dataset. To distinguish population-level trends from donor-specific relationships, these analyses were performed separately under vehicle and DEX, both across pooled donor cell strains and for each donor individually (Fig. 9; Supplemental Tables 1–2). Correlations were restricted to marker-positive cells as described in the Methods.

**Figure 9.**
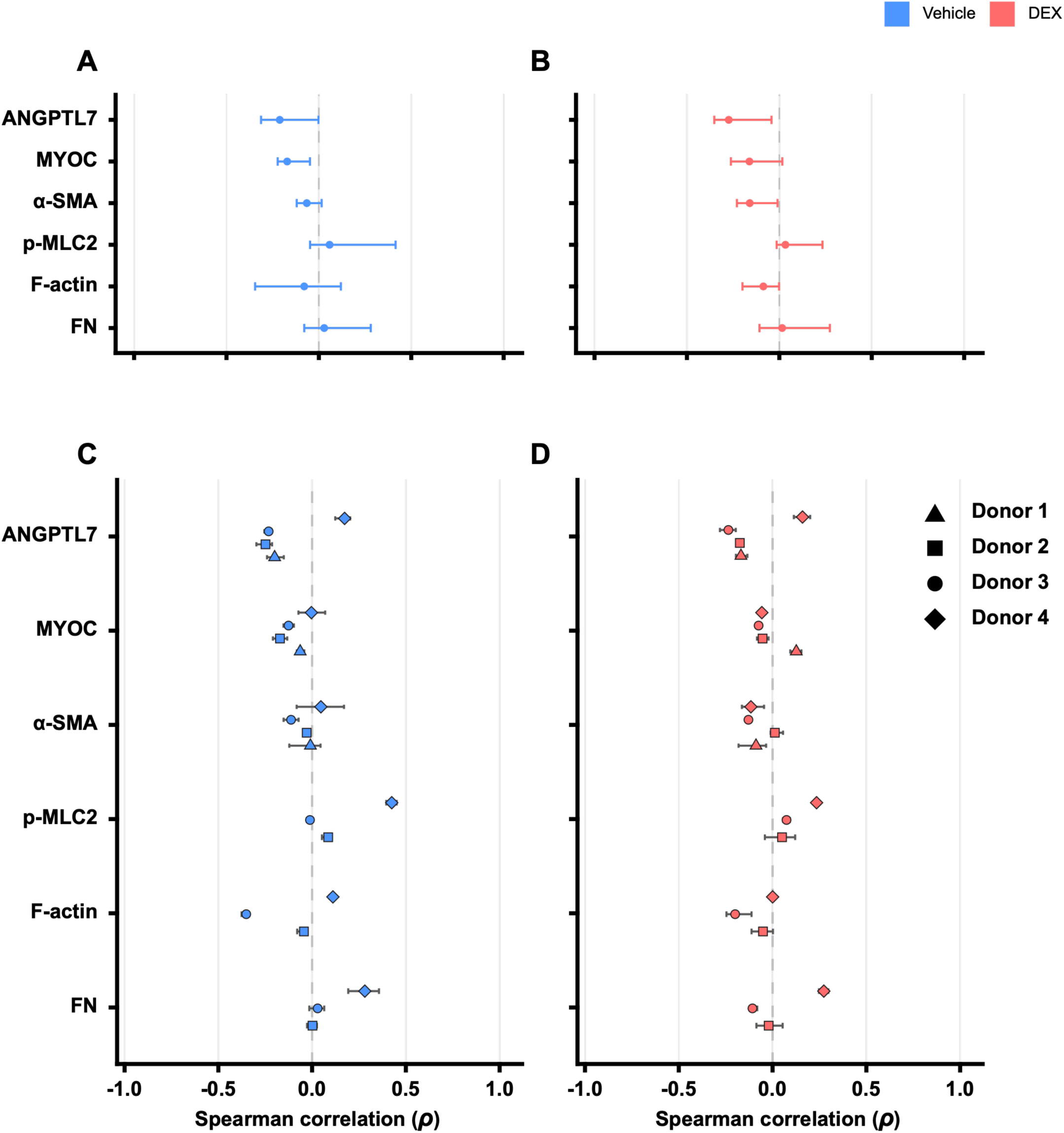
Individual marker abundance does not reliably predict single-cell contractile displacement. Single-cell correlations between contractile displacement and marker fluorescence intensity are shown for cells treated with vehicle (0.05% DMSO; **A**, **C**) or 100 nM DEX (**B**, **D**). Pooled correlations across donors are shown in **(A, B)** and donor-specific correlations in **(C, D)**. Points indicate Spearman correlation coefficients (rho; ρ), with horizontal bars indicating 95% confidence intervals (CIs). Measurements were collected at 6 h, corresponding to peak median contractile displacement in the live-cell assay. N = 4 human donor strains (TM349, TM403, TM404, TM405) for ANGPTL7, MYOC, and α-SMA and N = 3 human donor strains (TM403, TM404, TM405) for p-MLC2, F-actin, and fibronectin (FN), with a minimum of 2,000 cells analyzed per marker per condition. Only marker-positive cells were included in the analysis.

At the pooled level, correlations between protein abundance and contractile displacement were consistently weak across all six markers under both treatment conditions (|ρ| ≤ 0.274; Fig. 9A,B; Supplemental Table 1). Under vehicle, ANGPTL7 and MYOC exhibited the strongest negative associations (ρ = −0.212 and −0.171, respectively), whereas correlations for cytoskeletal and ECM-associated proteins were smaller and directionally uncertain. DEX modestly strengthened the negative association for ANGPTL7 (ρ = −0.274) but did not substantially alter the overall pattern of weak correlations across markers.

Despite these weak pooled associations, donor-specific analyses revealed considerable variation in the strength and direction of molecular and mechanical relationships (Fig. 9C,D; Supplemental Table 2). This was particularly evident for ANGPTL7, which exhibited consistently negative correlations in three donors under both treatments (ρ = −0.249 to −0.169), but positive correlations in a fourth donor (ρ = +0.173 under vehicle and +0.160 under DEX). MYOC showed a related pattern, with correlations remaining negative across donors under vehicle but becoming positive in one donor following DEX treatment. These opposing correlations reveal that the relationship between protein abundance and contractile displacement can differ across donors, even when the pooled population exhibits only weak associations.

Donor-specific variability was also evident among cytoskeletal and ECM-associated proteins, although the patterns differed by marker. For p-MLC2, one donor exhibited a moderate positive correlation under vehicle (ρ = +0.425), representing the strongest association observed across the entire analysis, whereas correlations in the remaining donors were substantially weaker. Similar donor-dependent differences were observed for FN and F-actin, with individual donors exhibiting stronger positive or negative associations than those suggested by the pooled estimates. In contrast, α-SMA correlations remained consistently weak across donors and treatments (|ρ| ≤ 0.129).

To determine whether the weak associations observed in the primary marker panel extended to additional TM-enriched proteins, we conducted an exploratory analysis of four recently described markers: fatty acid-binding protein 4 (FABP4), alpha-B crystallin (CRYAB), chitinase-3-like protein 1 (CHI3L1), and tomoregulin-2 (TMEFF2)^18,19^, using primary HTM cells from a single donor. Immunofluorescence staining confirmed expression of all four proteins in HTM cells adherent to collagen IV micropatterns (Supplementary Fig. 5C–F). Paired single-cell measurements revealed minimal correspondence between protein abundance and contractile displacement across all four markers, with cells exhibiting a broad range of displacement at similar protein abundance levels under both vehicle and DEX. Thus, the pattern of weak correlations extended to the TM-enriched proteins under both vehicle and DEX (|ρ| ≤ 0.21; Supplementary Fig. 5A,B).

Collectively, these findings demonstrate that intracellular protein abundance is only weakly associated with single-cell contractile displacement, despite the broad increase in contractile behavior induced by DEX. Furthermore, donor-specific analyses reveal that these weak pooled associations are not uniform, with relationships differing in both strength and direction across donors. Thus, individual marker abundance alone cannot explain the pronounced heterogeneity in HTM cell contractile behavior, suggesting that additional regulatory processes shape contractile output beyond what static protein abundance measurements capture.

## Discussion

In this study, we show that TM contractility is not a uniform property of the cell population but a dynamic, context-dependent single-cell mechanical state shaped by ECM identity, matrix stiffness, and pharmacologic perturbation. Using a high-throughput single-cell contractility assay, we show that ECM identity establishes distinct contractile programs, DEX selectively prolongs highly contractile states, ECM stiffness progressively constrains these responses, and ROCK inhibition collapses them into a uniformly low-contractility phenotype. Furthermore, static molecular marker abundance poorly predicts single-cell mechanical behavior, suggesting that the observed contractile phenotype reflects dynamic cellular regulation that molecular marker expression alone cannot explain.

### ECM Identity Establishes Distinct Contractile Programs

Our findings establish ECM identity as a primary determinant of intrinsic HTM mechanics, showing that ECM composition shapes contractile behavior without pharmacologic stimulation. Although previous collagen gel contraction assays have demonstrated that collagen substrates support HTM contractility, these approaches report population-averaged mechanical outputs and therefore cannot resolve how individual cells organize their mechanical behavior over time^20–22^. By measuring contractility at single-cell resolution, we found that collagen I, collagen IV, and fibronectin supported distinct mechanical phenotypes that differed not only in their contractile magnitude but also in their temporal evolution and population heterogeneity. Collagen IV was especially permissive for rapid and coordinated contractile activation, with the population converging toward an apparent shared contractile ceiling within the observation period. Collagen I engaged a related response more gradually and with incomplete population-wide convergence, whereas fibronectin maintained lower displacement and progressively increasing heterogeneity, with no evidence of convergence. Thus, ECM composition regulates not only the magnitude of HTM cell contractility but also the temporal organization of mechanical behavior across individual cells.

Fibrillar collagens such as collagen I are widely used to investigate cell contractility because of their load-bearing architecture and ability to support cell-mediated matrix deformation^23–25^. In contrast, collagen IV is generally viewed as a structural basement membrane component that contributes to filtration and tissue organization within the conventional outflow pathway, and its role in regulating cell mechanics has been comparatively understudied^5,26,27^. Our findings demonstrate that when presented in defined, mechanically permissive geometries, collagen IV can support robust HTM cell contractile behavior, suggesting a broader role for basement membrane components in TM biomechanics than previously appreciated.

In contrast, fibronectin did not simply support a weaker version of the response observed on collagens. Instead, it produced a qualitatively distinct mechanical phenotype characterized by persistently low displacement, increasing heterogeneity, and no evidence of convergence toward a shared contractile ceiling within the observation period. These findings are consistent with previous reports describing divergent, model-system-dependent effects of fibronectin on TM contractility and cytoskeletal organization^28–31^ and may reflect differences in integrin-mediated signaling and cytoskeletal regulation. Together, these findings demonstrate that ECM identity shapes not only contractile magnitude but also the kinetics, heterogeneity, and population-level organization of intrinsic HTM mechanical behavior.

### DEX Selectively Sustains Highly Contractile States

DEX-induced enhancement of TM contractility has been well documented in bulk contraction assays, including collagen gel and hydrogel models^20,21,32,33^. While these approaches demonstrate that glucocorticoids alter TM mechanics, they cannot resolve how the population-level response is distributed across individual cells. Specifically, they cannot distinguish whether DEX produces a uniform shift across the population, selectively enhances contractile behavior in an already responsive subset of cells, or both. Our single-cell analyses reveal that DEX selectively enhances and sustains the high-displacement upper tail of the contractile distribution rather than producing a uniform shift across the population.

Evidence for this selective DEX effect lies in the pre-existing contractile displacement distribution. Right-skewed displacement distributions were already present under basal conditions (Fig. 2), with similarly broad distributional patterns observed under vehicle and DEX treatment (Fig. 4). This selectivity extended into the relaxation phase, where the upper quartile remained elevated under DEX despite variable median trajectories (Fig. 5). The persistence of this high-contractility tail therefore emerged as the most reproducible relaxation-phase signature of DEX treatment. These findings indicate that glucocorticoid exposure does not create a new, uniformly hypercontractile state, but instead acts upon intrinsic mechanical heterogeneity by enhancing and sustaining a pre-existing high-contractility subpopulation.

This interpretation is consistent with a notion that HTM cells are morphologically, functionally, and transcriptionally diverse^13,18,19,34^. Our findings extend this framework along a mechanical axis by demonstrating that contractile behavior is itself dynamically regulated at the single-cell level. Whether this selective glucocorticoid effect on contractile behavior is associated with transcriptionally defined TM subpopulations remains unresolved, as the cells examined here were not transcriptionally profiled and human datasets have not yet converged on TM subtype number or identity^18,19^.

The selective nature of the DEX response also provides a mechanistic basis for the substantial donor-to-donor variability. Rather than imposing a uniform pharmacologic effect, DEX treatment appears to accentuate pre-existing differences in contractile capacity, with some donors exhibiting pronounced steroid effects and others more modest responses. Alternatively, apparent donor-specific mechanical phenotypes may partly reflect differences in the cellular composition of individual HTM isolates. Distinguishing donor-intrinsic differences from variation in cellular composition will require paired functional and molecular characterization of the same isolates. Consequently, population-averaged assays are likely to underestimate or obscure this structured heterogeneity, contributing to the variable and sometimes contradictory DEX effects reported previously^33,35^.

### ECM Stiffness Conditions Glucocorticoid Responsiveness

Our time-resolved analyses demonstrate that the defining mechanical effect of DEX is not accelerated contraction but prolonged maintenance of contractile activity. Under physiological stiffness, vehicle– and DEX-treated HTM cells reached similar peak contractile displacement rates during the initial contraction phase, indicating that glucocorticoid exposure does not substantially alter contraction initiation. Instead, DEX selectively prolonged contractile activity during the subsequent deceleration phase, delaying disengagement from contraction. This kinetic signature persisted into the relaxation phase, where DEX consistently maintained the highly contractile upper tail despite increasing divergence in donor-specific median trajectories.

This interpretation is consistent with previous studies showing that DEX promotes stress fiber formation, focal adhesion maturation, and expression of proteins involved in actomyosin organization and cell– ECM adhesion^36,37,38,39,40^. Our data extend these observations by providing direct functional evidence that glucocorticoids prolong active contractile engagement at the level of individual cells, and that this response is highly dependent on the surrounding mechanical environment^41^. As ECM stiffness increased, DEX progressively lost its ability to sustain prolonged contractile activity, and both the magnitude and heterogeneity of the steroid response became increasingly restricted. On substrates that approximate physiological stiffness, DEX prolonged contraction in the most mechanically active cells while preserving broad single-cell heterogeneity. Intermediate stiffness substantially attenuated this response, eliminating the prolonged deceleration phase while retaining a modest but reproducible increase in overall contractility. On pathologically stiff substrates, however, contractile behavior was almost completely abolished, and DEX produced no measurable mechanical response. These findings indicate that glucocorticoid responsiveness is not an intrinsic property of HTM cells but is strongly conditioned by the mechanical environment in which those cells reside. Progressive ECM stiffening is a hallmark of glaucomatous TM remodeling and is widely believed to contribute to increased aqueous humor outflow resistance^42,43,44^. In this context, our findings that increasing substrate stiffness was associated with progressively smaller and narrower distributions of single-cell contractile displacement, accompanied by reduced separation between vehicle– and DEX-treated cells, suggest that ECM stiffness substantially influences the observable mechanical responses of individual TM cells and the ability to resolve DEX-associated differences.

### ROCK Inhibition Establishes a Distinct Low-Contractility Mechanical State

ROCK signaling is a central regulator of actomyosin tension and has emerged as a major therapeutic target for lowering intraocular pressure^45,46,47,48,49,50–53^. Previous studies have frequently interpreted the ability of ROCK inhibitors to suppress glucocorticoid-induced contractility as evidence that they restore TM cells toward a normal mechanical phenotype. However, these conclusions have largely relied on bulk or endpoint measurements that cannot resolve how contractile behavior is organized across individual cells.

Our single-cell analyses demonstrate a different interpretation. Rather than selectively reversing the DEX-induced mechanical phenotype, ROCK inhibition broadly suppressed TM cell contractility. Under both ROCK inhibitor alone and combined DEX treatment, the broad spectrum of contractile states observed under vehicle and DEX conditions collapsed into a narrow population of uniformly low-displacement cells. This profound reduction in both donor-to-donor variability and single-cell heterogeneity indicates that ROCK inhibition establishes a mechanically distinct state rather than restoring basal contractile behavior. Consequently, what appears as rescue in conventional population-averaged assays is more accurately interpreted as global suppression of dynamic contractility.

This distinction may have important therapeutic implications. Physiological regulation of aqueous humor outflow likely depends on dynamic modulation of contractile activity rather than constitutive suppression of cellular force generation. If normal TM function requires continual adaptation to fluctuating mechanical loads, then broad elimination of contractile dynamics may represent a fundamentally different mechanical state from physiological homeostasis. Whether more selective modulation of Rho/ROCK signaling can suppress pathological contractility while preserving normal dynamic mechanical behavior remains an important question for future investigation and may inform the development of next-generation therapies targeting the conventional outflow pathway.

### Mechanical State Cannot Be Predicted from Static Molecular Identity

The inability of static molecular markers to predict single-cell mechanical behavior represents a second major conceptual finding of this study. Although cytoskeletal, extracellular matrix, and glaucoma-associated proteins are frequently interpreted as indicators of TM contractility, we found that protein abundance alone provided little information about the functional mechanical state of individual cells. Across all markers examined, including F-actin, p-MLC2, α-SMA, fibronectin, MYOC, and ANGPTL7, cells expressing similar protein levels exhibited the full spectrum of contractile behaviors, indicating that static molecular abundance and dynamic mechanical behavior are only weakly coupled at the single-cell level.

Quantitative correlation analyses supported this conclusion. Pooled correlations between marker abundance and contractility were uniformly weak under both vehicle and DEX treatment. Donor-stratified analyses further revealed that these weak pooled relationships often resulted from averaging divergent donor-specific associations rather than a universally absent biological relationship. Nevertheless, even within individual donors, correlations remained modest, demonstrating that molecular abundance alone is insufficient to predict the contractile behavior of HTM cells.

These findings suggest that functional mechanical behavior emerges from integrating multiple regulatory processes, including cytoskeletal organization, cell–ECM adhesion, intracellular signaling, and extracellular mechanical context, rather than from the abundance of any single protein. Similar molecular-mechanical decoupling has been reported in other mechanically active systems, including cardiomyocytes and stem cells, where canonical molecular markers incompletely predict functional behavior^54,55,56^. Our findings extend this principle to the trabecular meshwork and support the concept that dynamic mechanical state represents a complementary dimension of cellular identity that cannot be inferred from static molecular profiling alone.

### Limitations

Our study has several limitations. First, the native trabecular meshwork ECM is a complex composite of collagens, laminins, fibronectin, proteoglycans, and numerous matrix-associated proteins that collectively regulate cell behavior. Here, we intentionally adopted a reductionist approach by presenting defined ECM ligands on engineered micropatterns, allowing the independent contributions of ECM identity and stiffness to be examined without the confounding influence of overlapping biochemical cues. Although this simplified system does not fully recapitulate the native outflow pathway, it enabled direct interrogation of ligand-specific mechanical programs at single-cell resolution.

Second, experiments were performed on two-dimensional micropatterned substrates that do not reproduce the full three-dimensional architecture, fluid shear stress, pulsatile loading, or cyclic mechanical strain experienced by TM cells in vivo. These biomechanical stimuli undoubtedly contribute to conventional outflow physiology and may further influence contractile state. Nevertheless, the simplified geometry of the present platform provided precise control of matrix composition and mechanics while enabling high-throughput, time-resolved measurements of thousands of individual cells, capabilities that remain difficult to achieve in more complex experimental systems. Future integration of controlled flow, cyclic stretch, or three-dimensional ECM architectures with single-cell mechanical measurements should provide an increasingly physiologic view of TM mechanobiology.

Third, ROCK signaling was investigated using a single pharmacologic inhibitor at a concentration widely employed in TM mechanobiology studies. Although this facilitates comparison with previous work, off-target kinase inhibition cannot be excluded. Whether lower concentrations, alternative ROCK inhibitors, or pathway-selective modulation preserve physiological contractile dynamics while suppressing pathological mechanical states remains an important question for future investigation.

Finally, molecular and mechanical measurements were acquired at defined experimental endpoints and therefore cannot resolve potential temporal delays between signaling activation, protein accumulation, and functional contractile output. Because both molecular expression and mechanical behavior are themselves regulated by extracellular context, future studies integrating longitudinal single-cell mechanical measurements with transcriptomic or proteomic profiling of the same cells will be essential for defining the regulatory networks that establish and maintain distinct mechanical states.

Despite these limitations, the present study provides a framework for investigating TM mechanobiology at the level of individual cells. By resolving contractility dynamically rather than as a population average, we demonstrate that HTM contractility is organized as a spectrum of context-dependent mechanical states whose emergence is governed by ECM composition, progressively constrained by matrix stiffness, selectively prolonged by glucocorticoids, and globally suppressed by ROCK inhibition. Equally important, we show that these functional mechanical states cannot be reliably inferred from static molecular marker abundance alone, highlighting the importance of directly measuring cellular mechanics alongside molecular identity.

More broadly, these findings suggest that mechanical state represents a previously underappreciated dimension of TM cell identity. Resolving these dynamic states at single-cell resolution provides a framework for understanding how extracellular environment, pharmacologic perturbation, and inter-individual variability collectively regulate aqueous humor outflow in health and glaucoma. More broadly, this approach may apply to other mechanically regulated tissues in which population-averaged measurements obscure functionally important cellular heterogeneity.

