## Supplemental Information for "Dynamic regulation of single-cell contractile heterogeneity in the human trabecular meshwork"

*Corresponding

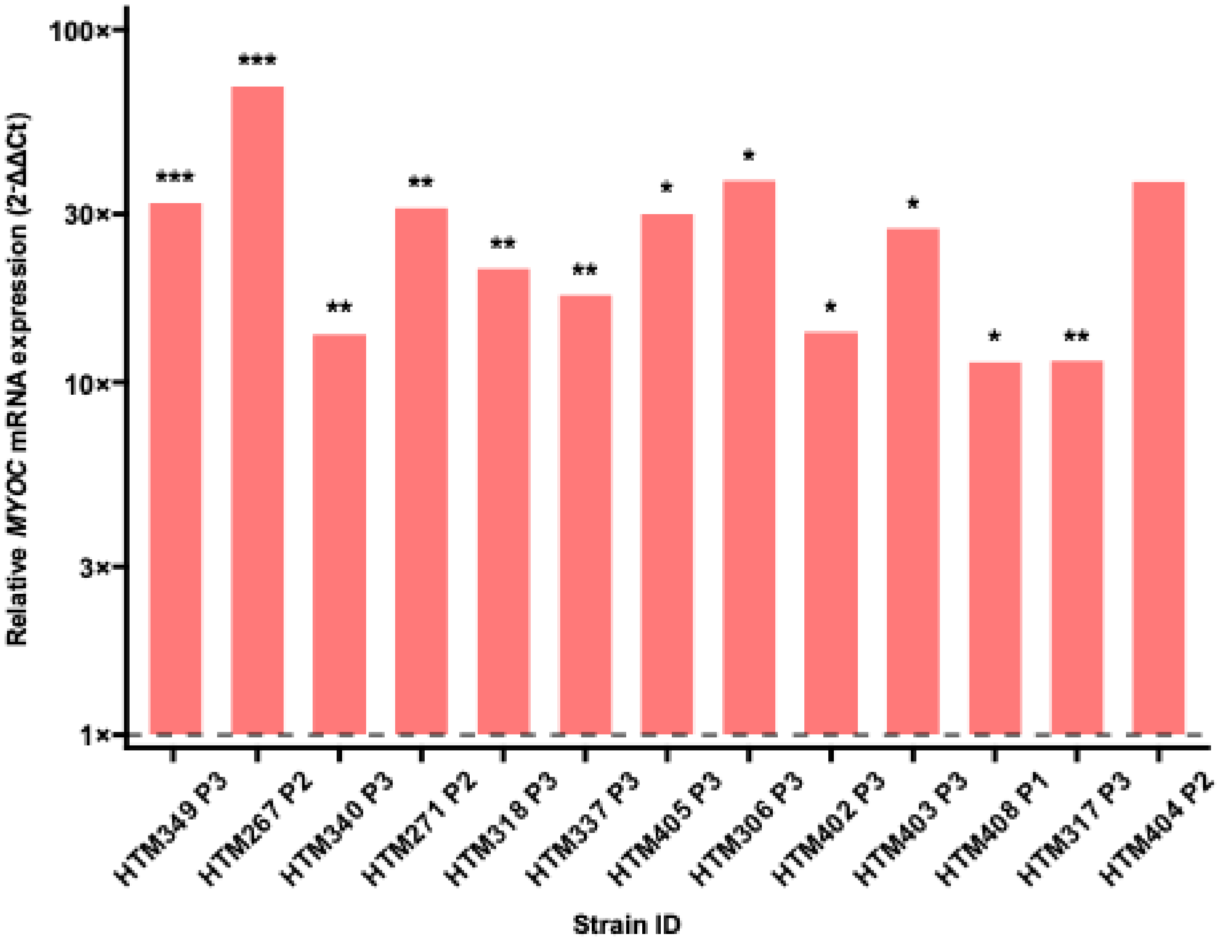


**Supplementary Figure 1. DEX-induced MYOC expression confirms HTM cell identity.** HTM cells (passage 1-3) were treated for 5 days with either vehicle control (0.05% DMSO) or 100 nM DEX. MYOC mRNA expression was quantified by TaqMan qRT-PCR, normalized to GAPDH, and analyzed using the ΔΔCt method. Bars show mean donor-level fold-change (2-ΔΔCt) relative to vehicle-treated controls. Statistical significance was assessed using a one-sided Welch's t-test with Holm correction across donors (*p < 0.05, **p < 0.01, ***p < 0.001). No significance test was performed for the donor with a single biological replicate.

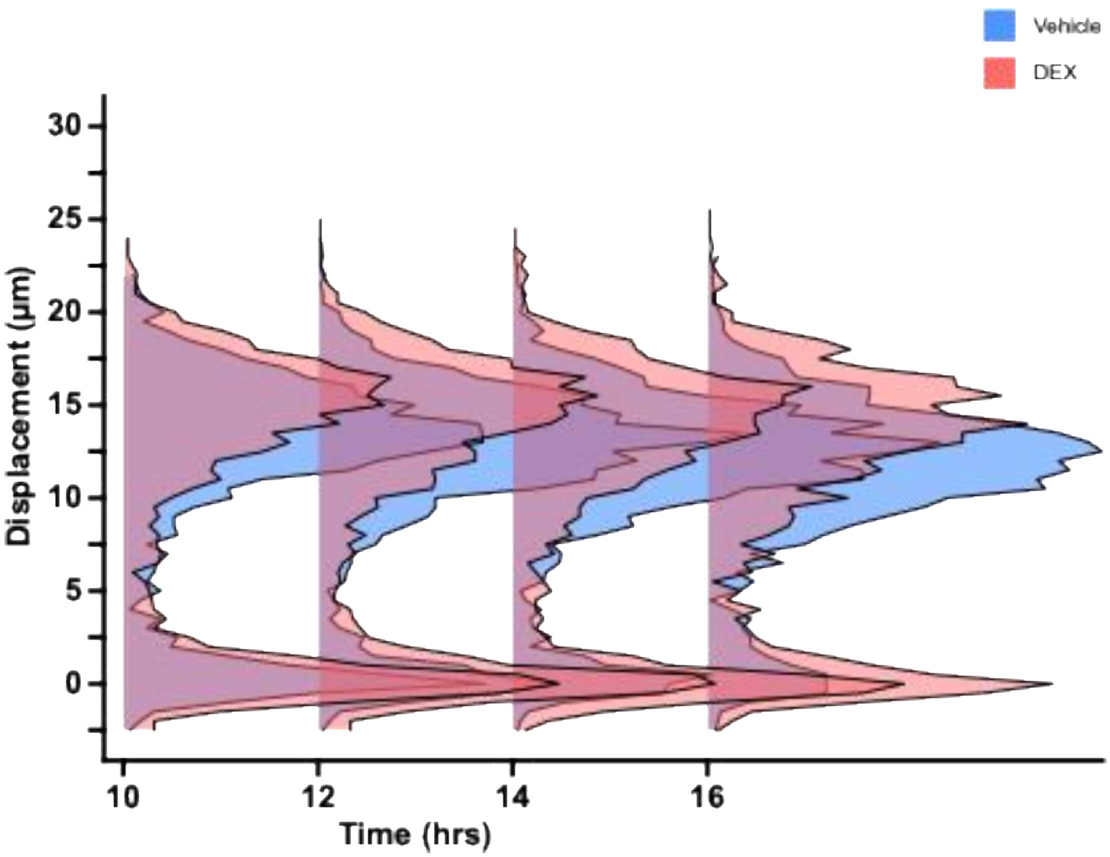


**Supplementary Figure 2. Single-cell displacement distributions for additional donor TM317 of the extended observation window.** Single-cell contractile displacement distributions at 10, 12, 14, & 16 hours on collagen IV micropatterned substrates of normal (4-8 kPa) stiffness. Cells were pre-treated for 5 days with either vehicle (0.05% DMSO) or 100 nM DEX. A minimum of 2,000 cells were analyzed per timepoint per condition.

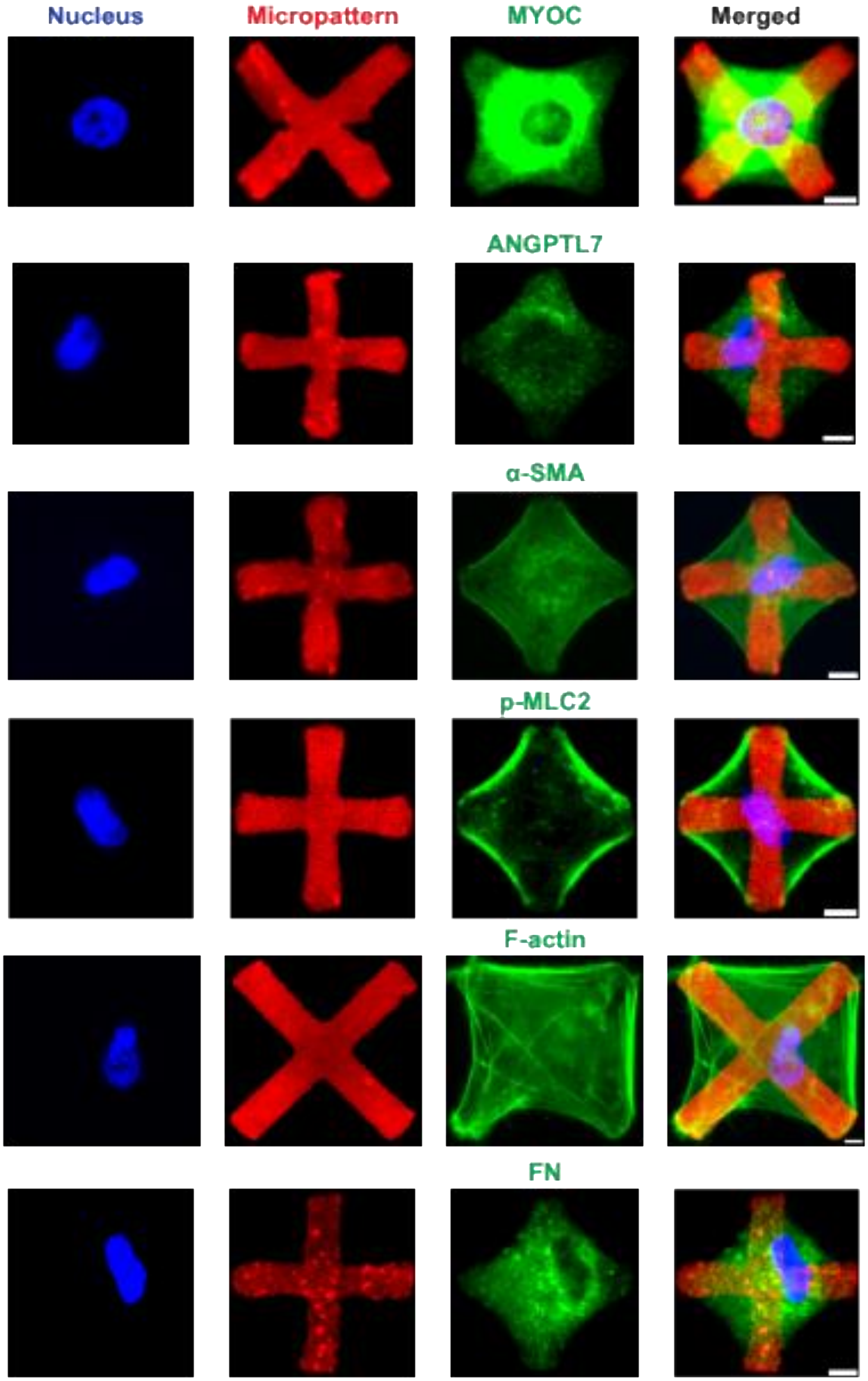


**Supplementary Figure 3. Representative immunofluorescence images showing protein distribution on collagen IV micropatterned substrates.** Images are shown for MYOC, ANGPTL7, α-SMA, p-MLC2, F-actin, and fibronectin (FN), with individual fluorescence channels and corresponding merged images. Collagen IV micropatterns are shown in red, marker proteins in green, and nuclei in blue. Scale bar = 10 μm.

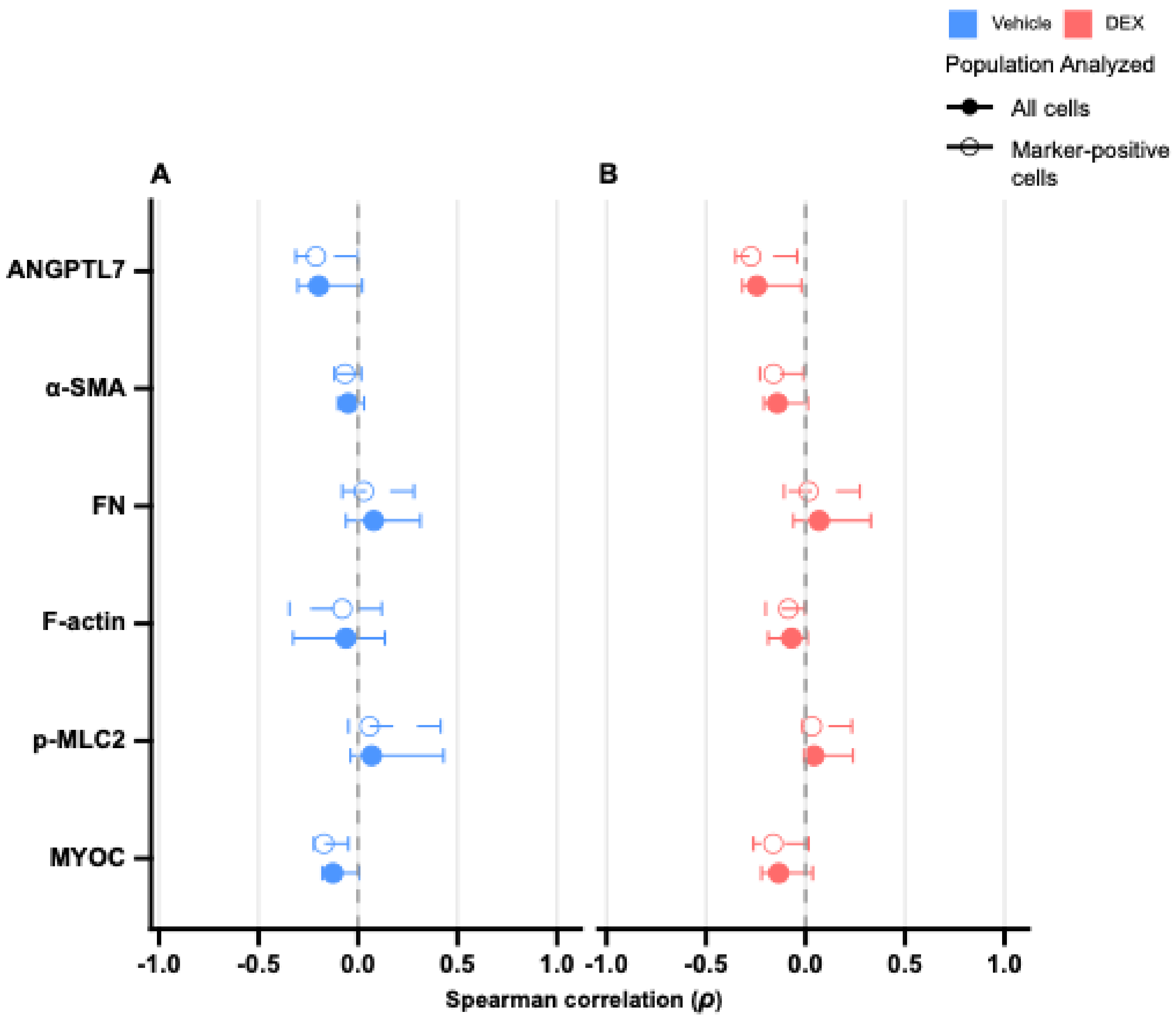


**Supplementary Figure 4. Influence of marker-negative cell inclusion on pooled Spearman correlation estimates.** Single-cell correlations between contractile displacement and marker fluorescence intensity under **(A)** vehicle (0.05% DMSO) or **(B)** 100 nM DEX. Each point represents a Spearman correlation coefficient ($\rho$), with horizontal bars indicating the 95% bootstrap confidence interval. Filled circles represent correlations computed across all cells regardless of marker expression status; open circles represent correlations restricted to marker-positive cells only. Data were collected at the 6-hour timepoint, corresponding to peak median contractile displacement in the live-cell assay. Data represent N = 4 donors (TM349, TM403, TM404, TM405) for ANGPTL7, MYOC, and α-SMA, and N = 3 donors (TM403, TM404, TM405) for p-MLC2, F-actin, and fibronectin (FN), with a minimum of 2,000 cells analyzed per marker per condition.

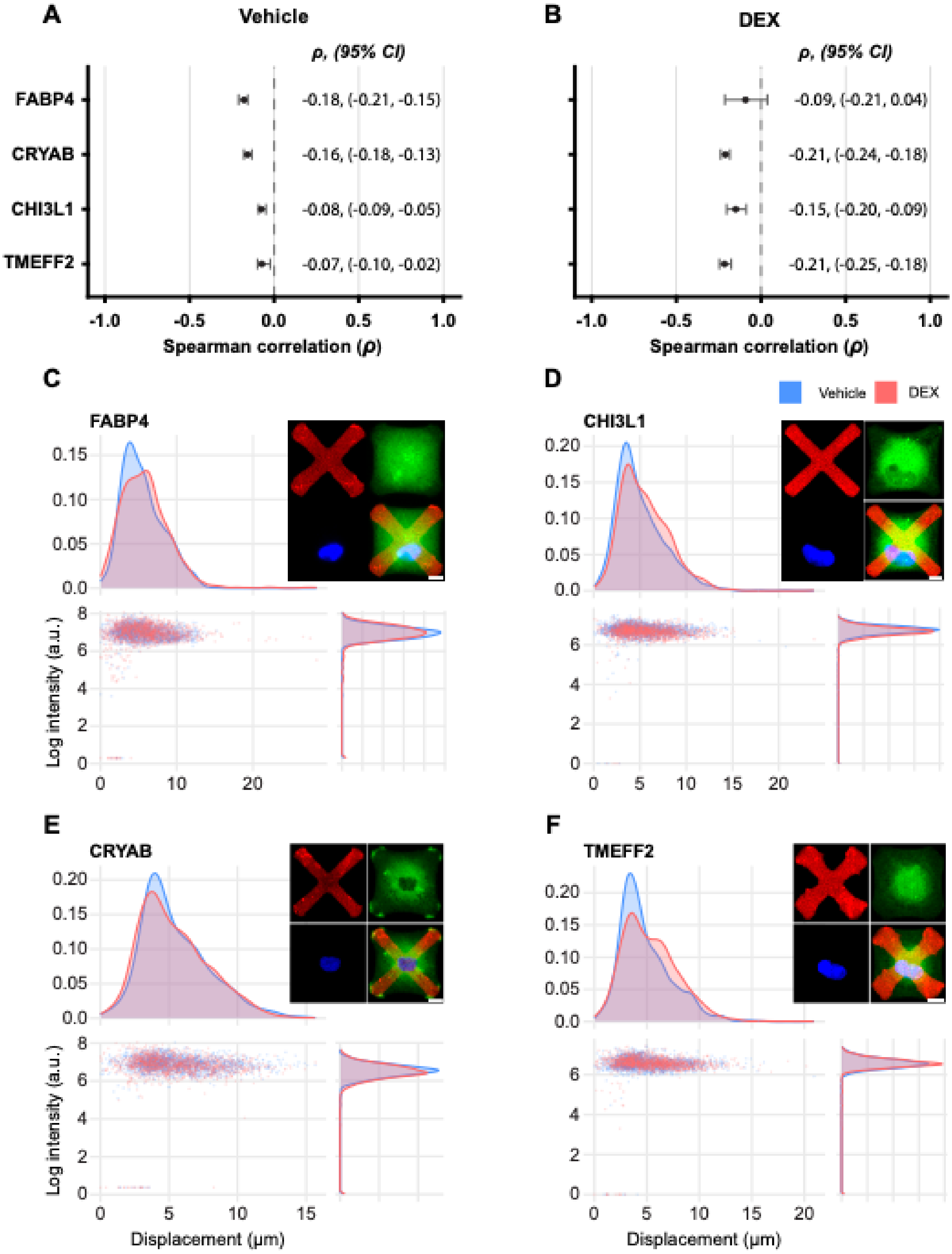

A
Vehicle
ρ, (95% CI)
FABP4
CRYAB
CHI3L1
TMEFF2
-0.18, (-0.21, -0.15)
-0.16, (-0.18, -0.13)
-0.08, (-0.09, -0.05)
-0.07, (-0.10, -0.02)
-1.0
-0.5
0.0
0.5
1.0
Spearman correlation (ρ)
B
DEX
ρ, (95% CI)
-0.09, (-0.21, 0.04)
-0.21, (-0.24, -0.18)
-0.15, (-0.20, -0.09)
-0.21, (-0.25, -0.18)
Spearman correlation (ρ)
C
FABP4
D
Vehicle
DEX
CHI3L1
Log intensity (a.u.)
E
CRYAB
F
TMEFF2
Displacement (μm)
Displacement (μm)

**Supplementary Figure 5. Single-cell correlations between displacement and additional TM-enriched protein markers.** (A–B) Spearman correlations ($\rho$) between single-cell contractile displacement and protein abundance for FABP4, CRYAB, CHI3L1, and TMEFF2 under vehicle control and DEX treatment. HTM cells were fixed, permeabilized, and immunostained 6 h after seeding on collagen IV micropatterned substrates (4–8 kPa). (C–F) Single-cell scatter plots with marginal distributions illustrating the relationship between contractile displacement and log-transformed protein intensity for (C) FABP4, (D) CHI3L1, (E) CRYAB, and (F) TMEFF2. Blue and red indicate vehicle and DEX conditions, respectively. Insets show representative immunocytochemistry images of HTM cells stained for the indicated marker (green), micropattern (red), and nuclei (blue). Data were obtained from a single primary HTM donor (TM349), with a minimum of 1,500 cells analyzed per marker per condition.

**Supplementary Table 1. Donor characteristics for primary HTM cell strains.**

| Strain ID | Age | Sex | Race | Passage |
|---|---|---|---|---|
| HTM267 | 42 | Male | Caucasian | 2 |
| HTM271 | 45 | Female | n.d. | 2 |
| HTM306 | 18 | Female | n.d. | 3 |
| HTM318 | 15 | Female | Caucasian | 3 |
| HTM337 | 19 | n.d. | n.d. | 3 |
| HTM340 | 35 | n.d. | n.d. | 3 |
| HTM349 | 38 | Male | black | 3 |
| HTM402 | 70 | Female | n.d. | 3 |
| HTM403 | 68 | n.d. | n.d. | 3 |
| HTM404 | 32 | Female | Hispanic | 2 |
| HTM405 | 62 | n.d. | n.d. | 3 |
| HTM408 | n.d. | n.d. | n.d. | 1 |
| HTM317 | 23 | Male | n.d | 3 |

Donor age, sex, and race were obtained from tissue provider records when available. Passage number refers to the culture passage used for the experiments. “n.d.” denotes information not disclosed by the tissue donor.

**Supplementary Table 2. Effect of Marker-Negative Cell Inclusion on Pooled Spearman Correlation Estimates**

| Marker | Treatment | Set | Effect | | |
|---|---|---|---|---|---|
| | | | ρ | CI Lower | CI Upper |
| ANGPTL7 | VEH | Marker-positive cells | -0.212 | -0.313 | -0.002 |
| ANGPTL7 | VEH | All cells | -0.198 | -0.302 | 0.021 |
| MYOC | VEH | Marker-positive cells | -0.171 | -0.223 | -0.048 |
| MYOC | VEH | All cells | -0.126 | -0.180 | 0.007 |
| α-SMA | VEH | Marker-positive cells | -0.065 | -0.120 | 0.015 |
| α-SMA | VEH | All cells | -0.049 | -0.102 | 0.030 |
| p-MLC2 | VEH | Marker-positive cells | 0.058 | -0.048 | 0.415 |
| p-MLC2 | VEH | All cells | 0.068 | -0.038 | 0.430 |
| F-actin | VEH | Marker-positive cells | -0.079 | -0.345 | 0.119 |
| F-actin | VEH | All cells | -0.061 | -0.326 | 0.135 |
| FN | VEH | Marker-positive cells | 0.029 | -0.079 | 0.281 |
| FN | VEH | All cells | 0.079 | -0.060 | 0.314 |
| ANGPTL7 | DEX | Marker-positive cells | -0.274 | -0.352 | -0.042 |
| ANGPTL7 | DEX | All cells | -0.244 | -0.318 | -0.021 |
| MYOC | DEX | Marker-positive cells | -0.162 | -0.262 | 0.016 |
| MYOC | DEX | All cells | -0.134 | -0.221 | 0.036 |
| α-SMA | DEX | Marker-positive cells | -0.160 | -0.229 | -0.010 |
| α-SMA | DEX | All cells | -0.141 | -0.210 | 0.011 |
| p-MLC2 | DEX | Marker-positive cells | 0.033 | -0.014 | 0.233 |
| p-MLC2 | DEX | All cells | 0.043 | -0.008 | 0.239 |
| F-actin | DEX | Marker-positive cells | -0.086 | -0.200 | -0.002 |
| F-actin | DEX | All cells | -0.070 | -0.187 | 0.010 |
| FN | DEX | Marker-positive cells | 0.015 | -0.107 | 0.273 |
| FN | DEX | All cells | 0.069 | -0.061 | 0.328 |

Pooled-level Spearman correlation coefficients ($\rho$) between single-cell marker fluorescence intensity and contractile displacement, computed separately within vehicle- and DEX-treated cell populations. 95% confidence interval (CI) lower and upper bounds were derived by bootstrap resampling. Correlations are reported for analyses including all cells regardless of marker expression status and for analyses restricted to marker-positive cells only, to illustrate the influence of marker-negative cell inclusion on correlation estimates. Graphical summaries are shown in Supplementary Figure #.

**Supplementary Table 3. Statistical summary of Spearman correlations at pooled level**

| Marker | Treatment | Effect | | |
|---|---|---|---|---|
| | | ρ | CI Lower | CI Upper |
| ANGPTL7 | VEH | -0.212 | -0.313 | -0.002 |
| MYOC | VEH | -0.171 | -0.224 | -0.050 |
| α-SMA | VEH | -0.065 | -0.120 | 0.015 |
| p-MLC2 | VEH | 0.058 | -0.048 | 0.425 |
| F-actin | VEH | -0.079 | -0.345 | 0.119 |
| FN | VEH | 0.029 | -0.079 | 0.281 |
| ANGPTL7 | DEX | -0.274 | -0.351 | -0.042 |
| MYOC | DEX | -0.162 | -0.259 | 0.013 |
| α-SMA | DEX | -0.160 | -0.229 | -0.007 |
| p-MLC2 | DEX | 0.033 | -0.014 | 0.235 |
| F-actin | DEX | -0.086 | -0.200 | 0.000 |
| FN | DEX | 0.015 | -0.107 | 0.265 |

Pooled-level Spearman correlation coefficients ($\rho$) between single-cell marker fluorescence intensity and contractile displacement, computed from data pooled across donors and computed separately under vehicle and DEX treatment conditions. 95% confidence interval (CI) lower and upper bounds were derived by bootstrap resampling. All analyses were restricted to marker-positive cells. Corresponding graphical summaries are shown in Figure 8A, B.

**Supplementary Table 4. Statistical summary of Spearman correlations at the donor-level.**

| | | | Effect | | |
|---|---|---|---|---|---|
| Donor | Marker | Treatment | ρ | CI Lower | CI Upper |
| TM349P3 | ANGPTL7 | VEH | -0.200 | -0.240 | -0.151 |
| TM403P3 | ANGPTL7 | VEH | -0.249 | -0.297 | -0.214 |
| TM404P2 | ANGPTL7 | VEH | -0.232 | -0.254 | -0.222 |
| TM405P3 | ANGPTL7 | VEH | 0.173 | 0.123 | 0.203 |
| TM349P3 | MYOC | VEH | -0.064 | -0.082 | -0.042 |
| TM403P3 | MYOC | VEH | -0.171 | -0.209 | -0.133 |
| TM404P2 | MYOC | VEH | -0.126 | -0.152 | -0.097 |
| TM405P3 | MYOC | VEH | -0.004 | -0.073 | 0.070 |
| TM349P3 | α-SMA | VEH | -0.009 | -0.122 | 0.045 |
| TM403P3 | α-SMA | VEH | -0.029 | -0.029 | -0.011 |
| TM404P2 | α-SMA | VEH | -0.112 | -0.152 | -0.072 |
| TM405P3 | α-SMA | VEH | 0.047 | -0.082 | 0.170 |
| TM403P3 | p-MLC2 | VEH | 0.086 | 0.053 | 0.091 |
| TM404P2 | p-MLC2 | VEH | -0.011 | -0.029 | 0.006 |
| TM405P3 | p-MLC2 | VEH | 0.425 | 0.396 | 0.451 |
| TM403P3 | F-actin | VEH | -0.044 | -0.080 | -0.044 |
| TM404P2 | F-actin | VEH | -0.351 | -0.376 | -0.336 |
| TM405P3 | F-actin | VEH | 0.111 | 0.093 | 0.126 |
| TM403P3 | FN | VEH | 0.002 | -0.027 | 0.026 |
| TM404P2 | FN | VEH | 0.030 | -0.015 | 0.065 |
| TM405P3 | FN | VEH | 0.281 | 0.193 | 0.357 |
| TM349P3 | ANGPTL7 | DEX | -0.169 | -0.195 | -0.134 |
| TM403P3 | ANGPTL7 | DEX | -0.174 | -0.181 | -0.162 |
| TM404P2 | ANGPTL7 | DEX | -0.236 | -0.281 | -0.196 |
| TM405P3 | ANGPTL7 | DEX | 0.160 | 0.113 | 0.200 |
| TM349P3 | MYOC | DEX | 0.126 | 0.094 | 0.154 |
| TM403P3 | MYOC | DEX | -0.053 | -0.083 | -0.021 |
| TM404P2 | MYOC | DEX | -0.074 | -0.083 | -0.063 |
| TM405P3 | MYOC | DEX | -0.057 | -0.072 | -0.036 |
| TM349P3 | α-SMA | DEX | -0.088 | -0.181 | -0.034 |
| TM403P3 | α-SMA | DEX | 0.012 | -0.010 | 0.056 |
| TM404P2 | α-SMA | DEX | -0.129 | -0.130 | -0.128 |
| TM405P3 | α-SMA | DEX | -0.116 | -0.163 | -0.045 |
| TM403P3 | p-MLC2 | DEX | 0.050 | -0.041 | 0.120 |
| TM404P2 | p-MLC2 | DEX | 0.074 | 0.056 | 0.093 |
| TM405P3 | p-MLC2 | DEX | 0.235 | 0.229 | 0.235 |
| TM403P3 | F-actin | DEX | -0.051 | -0.112 | 0.002 |
| TM404P2 | F-actin | DEX | -0.200 | -0.246 | -0.112 |
| TM405P3 | F-actin | DEX | 0.000 | -0.007 | 0.003 |
| TM403P3 | FN | DEX | -0.021 | -0.086 | 0.054 |
| TM404P2 | FN | DEX | -0.107 | -0.121 | -0.082 |
| TM405P3 | FN | DEX | 0.273 | 0.247 | 0.297 |

Donor-stratified Spearman correlation coefficients (*ρ*) between single-cell marker fluorescence intensity and contractile displacement, computed separately within vehicle- and DEX-treated cell populations for each donor.

95% confidence interval (CI) lower and upper bounds were derived by bootstrap resampling. All analyses were restricted to marker-positive cells. Graphical summaries are shown in Figure 8C, D.
